# Exploring the role of *E. faecalis* Enterococcal Polysaccharide Antigen (EPA) and lipoproteins in evasion of phagocytosis

**DOI:** 10.1101/2024.06.20.599754

**Authors:** Joshua S Norwood, Jessica L Davis, Bartłomiej Salamaga, Charlotte E Moss, Simon Johnston, Philip M Elks, Endre Kiss-Toth, Stéphane Mesnage

## Abstract

*Enterococcus faecalis* is an opportunistic pathogen frequently causing nosocomial infections. The virulence of this organism is underpinned by its capacity to evade phagocytosis, allowing dissemination in the host. Immune evasion requires a surface polysaccharide produced by all enterococci, known as the Enterococcal Polysaccharide Antigen (EPA). EPA consists of a cell wall-anchored rhamnose backbone substituted by strain-specific polysaccharides called “decorations”, essential for the biological activity of this polymer. However, the structural determinants required for innate immune evasion remain unknown, partly due to a lack of suitable validated assays. Here, we describe a quantitative, *in vitro* assay to investigate how EPA decorations alter phagocytosis. Using the *E. faecalis* model strain OG1RF, we demonstrate that a mutant with a deletion of the locus encoding EPA decorations can be used as a platform strain to express heterologous decorations, thereby providing an experimental system to investigate the inhibition of phagocytosis by strain-specific decorations. We show that the aggregation of cells lacking decorations is increasing phagocytosis and that this process does not involve the recognition of lipoproteins by macrophages. Collectively, our work provides novel insights into innate immune evasion by enterococci and paves the way for further studies to explore the structure/function relationship of EPA decorations.

## Introduction

*Enterococcus faecalis* is a commensal bacterium found in the human digestive tract that can cause hospital- and community-acquired infections. In elderly patients, immunocompromised hosts or following antibiotic-induced dysbiosis, *E. faecalis* is often responsible for a wide variety of diseases including infective endocarditis and peritonitis, as well as infections at urinary catheter, and other surgical, sites (1). *E. faecalis* displays a high resistance to extracellular stressors including mild disinfectants (2) and antibiotics commonly used to treat bacterial infections such as cephalosporins (3). The formation of biofilms is also a common feature of *E. faecalis*, further reducing the effectiveness of antibiotic treatments (4). Multi-species biofilms are of particular concern since *E. faecalis* can augment the virulence of other bacteria (5) and serve as a reservoir for antimicrobial resistance genes, particularly resistance to last-resort antibiotics such as vancomycin (6).

*E. faecalis* produces several virulence factors that have been studied in detail, but the exact mechanism of how this bacterium causes infections remains poorly understood. Virulence factors are not exclusively found in clinical isolates, and disease-causing strains can also colonize healthy individuals (7). The use of zebrafish as an experimental model of infection revealed that the ability of *E. faecalis* to avoid uptake by innate immune cells (macrophages and neutrophils) is critical for pathogenesis (8).

*E. faecalis* cell envelope composition and dynamics play an important role in resistance against innate immune effectors. Approximately 40 % of *E. faecalis* clinical isolates produce a capsular polysaccharide (9), which masks opsonic C3 molecules from recognition by phagocytes (10). Meanwhile, there is evidence that non-opsonic phagocytosis is inhibited by enterococcal glycolipids (11,12). The efficiency of *E. faecalis* uptake is further reduced by the activity of the autolysin AtlA, which prevents the formation of long chains of enterococci which are more readily phagocytosed (13).

*E. faecalis* has also evolved mechanisms to survive innate immune effectors. Expression of aggregation substance, an envelope-localised adhesin, for example, facilitates entry into neutrophils (14) and increases intracellular survival (15).

The enterococcal polysaccharide antigen (EPA) is a cell envelope polymer produced by all enterococci that contributes to virulence (16). EPA consists of a well-conserved rhamnose backbone decorated with covalently bound strain-specific polysaccharides called “decorations” (17,18). The chromosomal *epa* locus is subdivided into a conserved and a variable (*epa_var*) region. These two loci encode the biosynthetic machineries for the rhamnose backbone and the decoration polymers, respectively (18). Deletion of genes within either region significantly attenuates virulence (8,19). Current research suggests that EPA helps to maintain cell envelope integrity, thus increasing resistance to antimicrobial peptides (19) and favouring intracellular survival (20). In addition, mutants lacking EPA decorations are avirulent in zebrafish and more susceptible to uptake by macrophages *in vivo* (19). The mechanisms by which EPA decorations inhibit phagocytosis remain unknown.

In this work, we describe a quantitative *in vitro* phagocytosis assay to investigate how E. faecalis cell surface components modulate phagocytosis. We provide the proof of concept that *E. faecalis* OG1RF with a complete deletion of the decoration locus can be used as a platform strain to investigate (i) the structure/function relationship of EPA by performing heterologous expression of strain-specific EPA decorations, and (ii) the recognition of cell envelope components by phagocytes in the absence of EPA decorations. Finally, we show that EPA decorations reduce phagocytosis by inhibiting the aggregation of enterococcal cells, thereby promoting dissemination in the host.

## Results

### Setting up an *in vitro* phagocytosis assay using IBMDMs

We sought to design an *in vitro* assay to quantitatively assess non-opsonic phagocytosis of *E. faecalis* without compounding effects from other immune processes. Immortalised bone marrow-derived macrophages (iBMDMs), originally derived from oncogenic mice (21), were utilised as model host phagocytes to measure the uptake of *E. faecalis* OG1RF derivatives constitutively expressing GFP (13). Following internalisation by iBMDMs, the number of intracellular bacteria was determined by proxy, measuring the green fluorescence intensity of individual iBMDM cells (Fig. S1) (22).

Before we compared the uptake of different strains, two critical conditions were optimised: incubation time and multiplicity of infection (MOI). First, OG1RF wild-type bacteria were incubated alongside iBMDMs for 1 hour or 3 hours at 37 °C. Both test groups of iBMDMs showed a significant increase in fluorescence as compared to the no bacteria control, indicating that iBMDMs were internalising bacteria (Fig. 1a). Fluorescence intensity associated with iBMDMs was much lower after a 1 hour incubation as compared to after 3 hours. Based on these results, a 1 hour incubation time was chosen for all future experiments, to enable the characterisation of mutants more readily uptaken. Next, wild-type bacteria were added to iBMDMs at an MOI of 0, 1, 5, 20, or 100 before co-incubation (1 hour, 37 °C). A dose-response was observed, in which iBMDM fluorescence increased with increasing MOI (Fig. 1b). An MOI of 5 was chosen for future experiments, again to allow for the identification of mutants which are more readily phagocytosed.

**Fig. 1:**
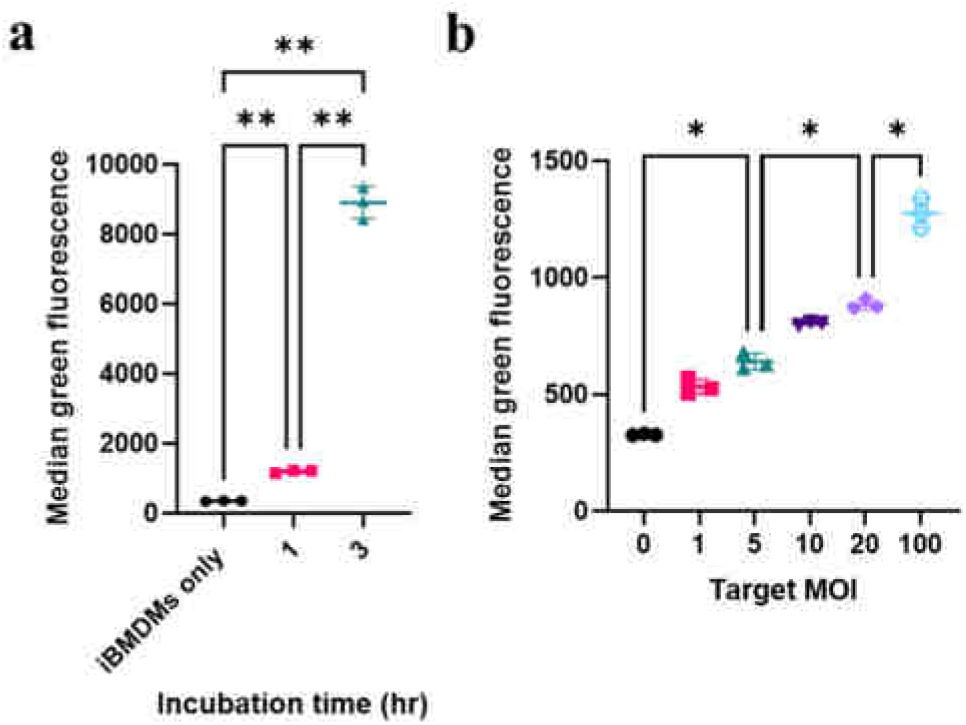
Setting up an assay to measure internalisation of GFP-labelled *E. faecalis* by iBMDMs. **(a)** Internalisation of *E. faecalis* after either 1 hour or 3 hours of incubation at 37 °C. The graph shows the average brightness of iBMDMs that contained bacteria (median green fluorescence in arbitrary units). To assess significance, a one-way ANOVA with Brown-Forsythe and Welch’s correction was performed, followed by Dunnett’s multiple comparisons test. *P*-values: iBMDMs only versus 1 hour, *P* = 0.0023; iBMDMs only versus 3 hours, *P* = 0.002; 1 hour versus 3 hours, *P* = 0.0025. **(b)** Internalisation (1 hour) of *E. faecalis* by iBMDMs according to bacterial dose. Again, statistical analysis was performed via a one-way ANOVA with Brown-Forsythe and Welch’s correction followed by Dunnett’s multiple comparisons test. *P*-values: MOI = 0 versus MOI = 5, *P* = 0.0188; MOI = 5 versus MOI = 20, *P* = 0.0132; MOI = 20 versus MOI = 100, *P* = 0.0137. For **(a)** and **(b)**, n = 3 technical replicates per condition, and error bars show mean values ± standard deviation (SD). *P*-value descriptors: *, *P* < 0.05; **, *P* < 0.01.

Another way of quantifying phagocytosis was to determine the percentage of macrophages that had taken up bacteria. When looking at this metric over increasing time/MOI, the same trends were observed (Fig. S2), supporting the previous conclusions and showing that it was not just a subpopulation of iBMDMs internalising bacteria. Finally, it was demonstrated that uptake was significantly higher at 37°C compared to 4°C (Fig. S3), indicating that *E. faecalis* uptake is an active process (23).

### *In vitro* uptake by iBMDMs to explore EPA structure/function

After optimising the conditions, the *in vitro* uptake assay was benchmarked using a mutant producing an EPA polysaccharide devoid of decorations (with a 17.6 kbp deletion of the *epa_var* region; strain Δ*epa_var*). As expected, the mutant displayed a significant increase in internalisation as compared to wild-type (Fig. 2a), confirming that EPA decorations facilitate escape from phagocytosis by macrophages. The mutant’s phenotype was fully complemented by a plasmid encoding the *epa_var* locus (Fig. 2a; pILvar_O). The empty vector pIL252 had no significant impact on phagocytosis, confirming that protection was due to OG1RF decorations. Interestingly, there was no difference in the percentage of iBMDMs harbouring bacteria between wild-type, Δ*epa_var* and complemented strains (Fig. S4a), but fluorescence intensity associated with iBMDMs significantly increased when mutant bacteria were administered (Fig. S4b). Our findings were verified by performing fluorescence microscopy analysis on iBMDMs incubated with wild-type, mutant, or complemented bacteria (Fig. S4c-d).

**Fig. 2:**
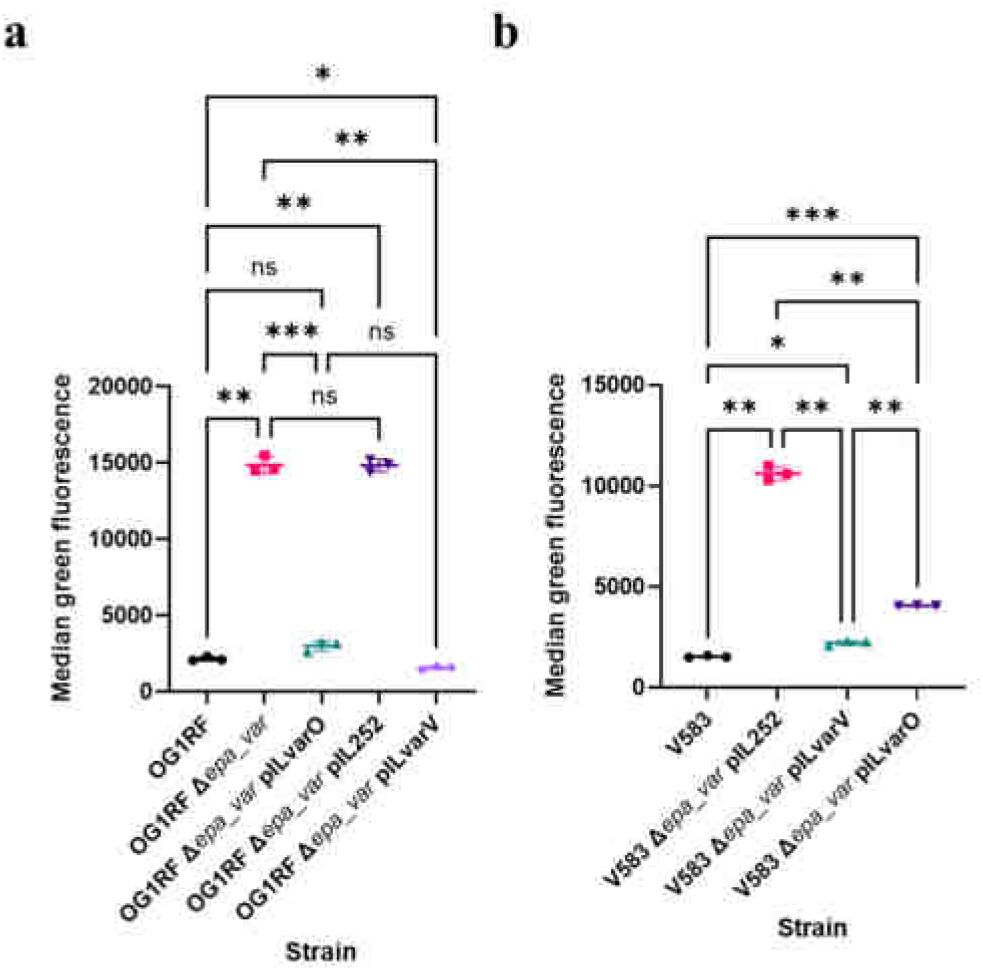
EPA decorations from strain V583 protect OG1RF Δ*epa_var* from phagocytosis, and vice versa. **(a)** Phagocytosis of OG1RF Δ*epa_var* transformed with an empty vector (pIL252) or a vector expressing V583 EPA decorations (pILvarV). Results are shown for one experiment with three technical replicates per group; these results are representative of three independent experiments. Statistical analysis was performed by one-way ANOVA with Brown-Forsythe and Welch’s correction followed by Dunnett’s multiple comparisons test. *P*-values: OG1RF versus Δ*epa_var*, *P* = 0.0025; OG1RF versus pILvarO, *P* = 0.118; OG1RF versus pIL252, *P* = 0.0014; OG1RF versus pILvarV, *P* = 0.0274; Δ*epa_var* versus pILvarO, *P* = 0.0003; Δ*epa_var* versus pIL252, *P* > 0.999; Δ*epa_var* versus pILvarV, *P* = 0.0023; pILvarO versus pILvarV, *P* = 0.0691. **(b)** Phagocytosis of V583 Δ*epa_var* transformed with pIL252 or a vector expressing OG1RF EPA decorations (pILvarO). Results are shown for one experiment with three technical replicates per group; these results are representative of three independent experiments. Statistical analysis was performed by one-way ANOVA with Brown-Forsythe and Welch’s correction followed by Dunnett’s multiple comparisons test. *P*-values: V583 versus pIL252, *P* = 0.0015; V583 versus pILvarV, *P* = 0.0115; V583 versus pILvarO, *P* = 0.0004; pIL252 versus pILvarV, *P* = 0.0020; pIL252 versus pILvarO, *P* = 0.0029; pILvarV versus pILvarO, *P* = 0.0047. Key to *P*-values: ns, not significant; *, *P* < 0.05; **, *P* < 0.01; ***, *P* < 0.001.

With the assay benchmarked, we sought to investigate if we could compare the function of strain-specific EPA decorations by doing cross-complementation experiments (24). As a proof of concept, we complemented the OG1RF Δ*epa_var* strain with a plasmid encoding the decoration from *E. faecalis* V583. Heterologous complementation revealed that V583 decorations offer a similar level of protection as OG1RF decorations (Fig. 2a). This was also observed in the inverse experiment (Fig. 2b), where heterologous expression of OG1RF decorations significantly reduced the uptake of V583 Δ*epa_var* (as compared to the empty vector). Altogether, these findings show that strain-specific EPA decorations can cross-complement one another, suggesting a conserved protective mechanism.

### Aggregation of the Δ*epa_var* mutant contributes to increased phagocytosis

The increase in median green fluorescence without an increase in the percentage GFP-positive macrophages (Fig. S4) suggested that more Δ*epa_var* cells are internalised as compared to wild type cells. A defect in bacterial daughter cell separation, leading to the formation of longer bacterial cell chains, has been suggested to increase bacterial uptake by phagocytes (13). We therefore decided to investigate if the morphology of the *epa_var* mutant cells is contributing to an increased phagocytosis. We started by comparing growth of the wild-type, mutant, and complemented strains. When grown in BHI broth at 37°C, OG1RF Δ*epa_var* displayed a significant increase in doubling time compared to wild-type (Fig. S5), indicating that EPA decorations help to maintain normal bacterial growth.

We noticed that the Δ*epa_var* mutant consistently showed fewer CFU counts versus wild-type cells when plated. To investigate this formally, serial dilutions of exponential cultures were plated and CFU counts were made and normalised to OD_600_ = 0.3. Δ*epa_var* exponential cultures consistently showed a decrease in CFU/ml compared to both the wild-type and the complemented strain (Fig. 3a). To investigate the phenotypes observed in more detail, exponential-phase Δ*epa_var* bacteria were analysed via confocal microscopy. Peptidoglycan cell wall shape and septum formation were visualised by labelling the bacteria with an Alexa555 NHS ester and a fluorescent D-amino acid (HADA), respectively (Fig. S6a). Mutant bacteria displayed single septa running perpendicular to the direction of cell division, suggesting that division was occurring normally. However, when compared to wild-type and complemented bacteria, Δ*epa_var* bacterial cells were significantly shorter in length and greater in width, giving them a more spherical appearance (Fig. S6b-d). In addition, microscopic analysis revealed more evidence that Δ*epa_var* bacteria form aggregates, with large, amorphous clumps of bacteria prevalent (Fig. 3b). In contrast, wild-type bacteria tended to be arranged as more discrete diplococci. To quantify the putative aggregation phenotype, FSC measurements were taken for wild-type, mutant, and complemented bacteria via flow cytometry (Fig. 3c). Δ*epa_var* bacteria displayed a significant increase in FSC, which is consistent with the formation of aggregates. Sonication of Δ*epa_var* bacteria decreased FSC and increased CFU count, which is also consistent with the bacterial cell aggregation hypothesis (Fig. 3d-e).

**Fig. 3:**
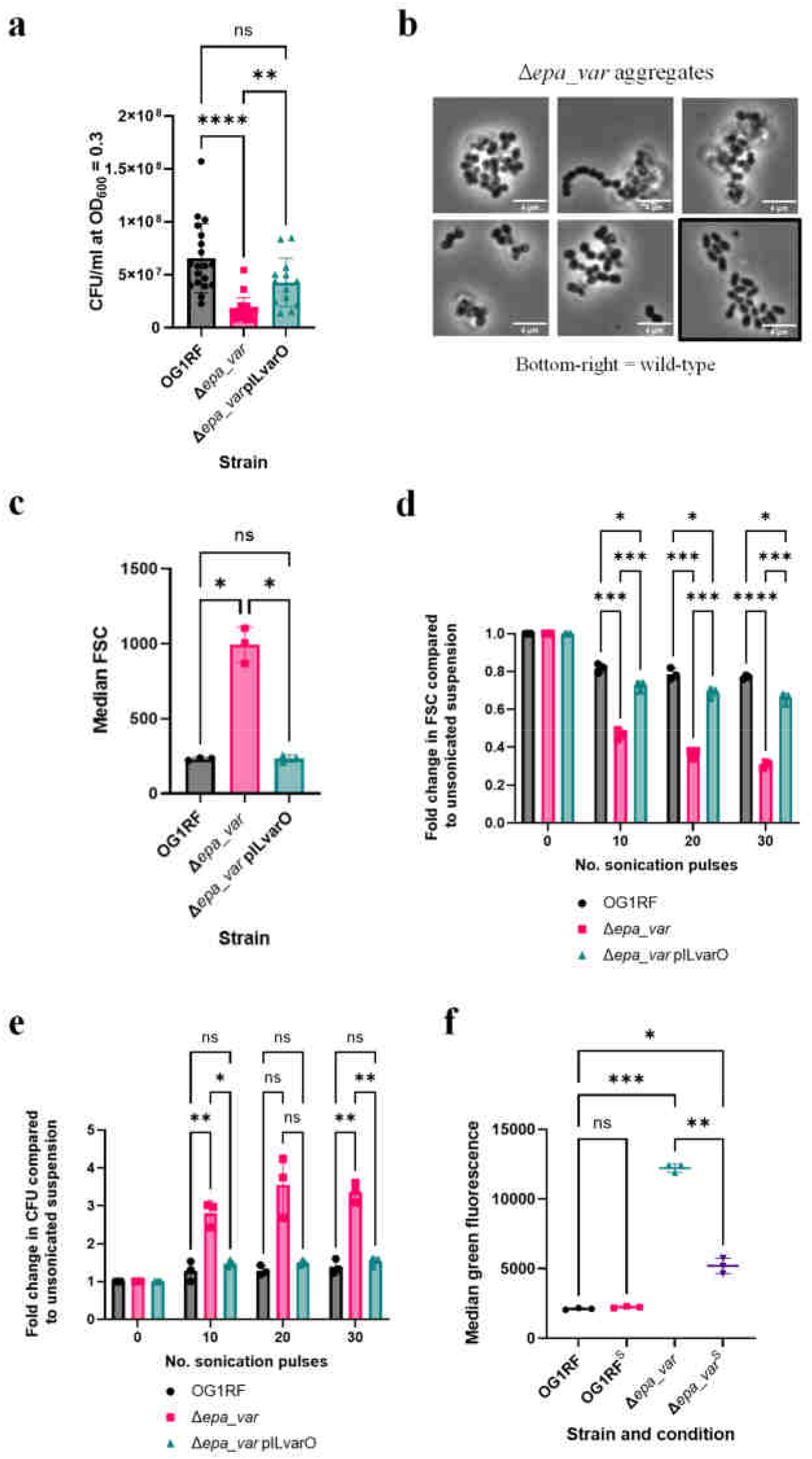
EPA decorations contribute to reduced internalisation by reducing bacterial cell aggregation. **(a)** CFU/ml of *E. faecalis* OG1RF Δ*epa_var* versus wild-type at OD600 = 0.3. Statistical analysis was performed by doing a one-way ANOVA with Brown-Forsythe and Welch’s correction, followed by Dunn’s multiple comparisons test. *P*-values: OG1RF versus Δ*epa_var*, *P* < 0.0001; OG1RF versus pILvarO, *P* = 0.0769; Δ*epa_var* versus pILvarO, *P* = 0.0062. Number of biological replicates per group: wild-type, n = 19; Δ*epa_var*, n = 20; Δ*epa_var* pILvarO, n = 13. **(b)** Representative phase contrast images of aggregates formed by *E. faecalis* OG1RF Δ*epa_var* bacteria. A representative image of wild-type bacteria (bottom right, boxed) is provided for comparison. **(c)** Median FSC of early exponential-phase OG1RF wild-type, Δ*epa_var*, or Δ*epa_var* complemented bacteria. For each group, n = 3 biological replicates. Statistical analysis was performed via a one-way ANOVA with Brown-Forsythe and Welch’s correction, followed by Dunnett’s multiple comparisons test. *P*-values: OG1RF versus Δ*epa_var*, *P* = 0.0167; OG1RF versus pILvarO, *P* = 0.996; Δ*epa_var* versus pILvarO, *P* = 0.0174. **(d)** Fold decrease in bacterial particle FSC compared to suspensions before sonication. Three biological replicates per group. Statistical analysis was performed by doing a two-way ANOVA followed by Tukey’s multiple comparisons test. No. pulses = 10: OG1RF versus Δ*epa_var*, *P* = 0.0002; OG1RF versus pILvarO, *P* = 0.0227; Δ*epa_var* versus pILvarO, *P* = 0.0004. No. pulses = 20: OG1RF versus Δ*epa_var*, P = 0.0002; OG1RF versus pILvarO, *P* = 0.0320; Δ*epa_var* versus pILvarO, *P* = 0.0002. No. pulses = 30: OG1RF versus Δ*epa_var*, *P* < 0.0001; OG1RF versus pILvarO, *P* = 0.0211; Δ*epa_var* versus pILvarO, *P* = 0.0004. **(e)** Fold increase in CFU count compared to bacterial suspensions before sonication. All counts were normalised to OD600 = 0.3. Three biological replicates per group. Statistical analysis was performed by doing a two-way ANOVA followed by Tukey’s multiple comparisons test. No. pulses = 10: OG1RF versus Δ*epa_var*, *P* = 0.0082; OG1RF versus pILvarO, *P* = 0.537; Δ*epa_var* versus pILvarO, *P* = 0.0278. No. pulses = 20: OG1RF versus Δ*epa_var*, *P* = 0.0651; OG1RF versus pILvarO, *P* = 0.162; Δ*epa_var* versus pILvarO, *P* = 0.0821. No. pulses = 30: OG1RF versus Δ*epa_var*, *P* = 0.0015; OG1RF versus pILvarO, *P* = 0.580; Δ*epa_var* versus pILvarO, *P* = 0.0043. **(f)** iBMDM-mediated phagocytosis of sonicated (^S^) or unsonicated bacteria. Sonicator settings = 20 pulses using 20% amplitude. In this experiment, three technical replicates were performed per group. Statistical analysis was performed using a one-way ANOVA with Brown-Forsythe and Welch’s correction followed by Dunnett’s multiple comparisons test. *P*-values: OG1RF versus OG1RF^S^, *P* = 0.247; OG1RF versus Δ*epa_var*, *P* = 0.0009; OG1RF versus Δ*epa_var*^S^, *P* = 0.0317; Δ*epa_var* versus Δ*epa_var*^S^, *P* = 0.0011. Key to *P*-values: ns, not significant; *, *P* < 0.05; **, *P* < 0.01; ***, *P* < 0.001; ****, *P* < 0.0001.

### Recognition of surface lipoproteins is not responsible for the increased phagocytosis in the absence of EPA decorations

The presence of EPA decorations at the cell surface prevents other cell envelope components from being recognised by immune receptors. In group B streptococci, the capsular polysaccharide masks a streptococcal lipoprotein from being recognised by macrophages by scavenger receptor A (25). enterococcal lipoproteins are known to activate pro-inflammatory signalling cascades (26) and may contribute to *E. faecalis*-associated intestinal inflammation (27). To test the role of lipoproteins in uptake, we generated *E. faecalis* OG1RF mutants with an in-frame deletion of *lgt (OG1RF_11459)* in both the wild-type (Fig. S7a) and Δ*epa_var* backgrounds (Fig. S7b). This gene encodes prolipoprotein diacylglyceryl transferase, the enzyme responsible for anchoring lipoproteins onto the enterococcal cell surface (28). A deletion of *lgt* in *E. faecalis* V583 led to an increase in lipoprotein shedding into the culture supernatant (29). We used a TCA-based precipitation method to purify proteins from culture supernatants and profile them via SDS-PAGE (Fig. 4a). More protein species were indeed detected in Δ*lgt* culture supernatants than were seen in parental ones. Furthermore, this phenotype could be complemented with an inducible expression system. Altogether, these findings suggest that our mutants lack Lgt activity.

**Fig. 4:**
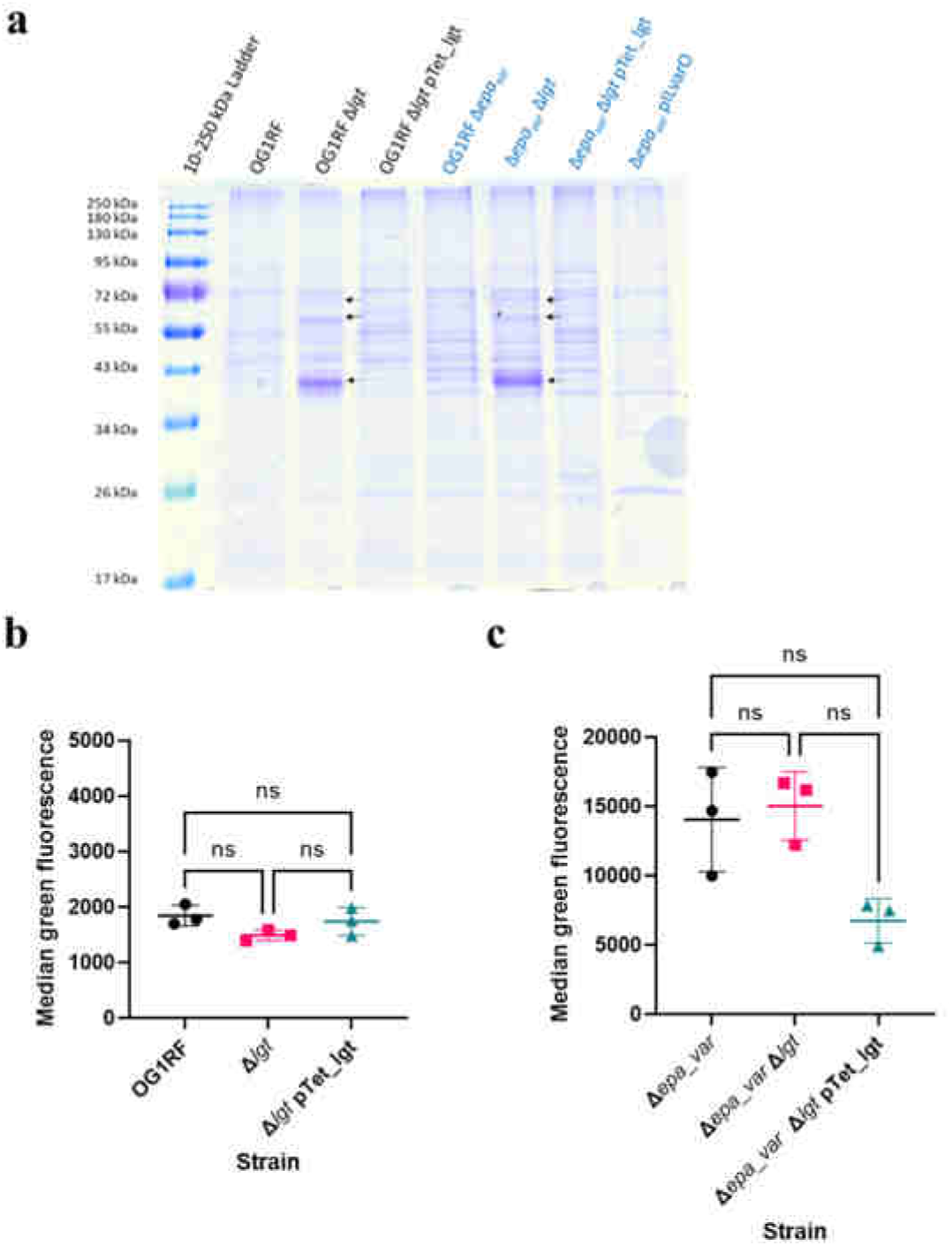
Characterisation of the OG1RF Δ*lgt* and Δ*epa_var* Δ*lgt* mutants. **(a)** Analysis of proteins released into the culture supernatant by *E. faecalis* cells in early exponential phase (OD600 ≈ 0.3). Deletion of *lgt* results in additional proteins shed (black arrows). **(b)** Phagocytosis of OG1RF Δ*lgt*. Statistical analysis was performed via one-way ANOVA with Brown-Forsythe and Welch’s corrections, followed by Dunnett’s multiple comparisons test (n = 3 biological replicates per group). **(c)** Phagocytosis of Δ*epa_var* Δ*lgt*. Same statistical analysis method as C (n = 3 biological replicates per group). *P*-values: ns, not significant; *, *P* < 0.05; **, *P* < 0.01; ***, *P* < 0.001; ****, *P* < 0.0001.

Next, we measured the uptake of these mutants by iMBDMs (Fig. 4b-c). Deletion of *lgt* did not lead to any significant change in phagocytosis, irrespective of the production of EPA decorations. This suggests that immune evasion is not due to EPA decorations masking lipoproteins.

## Discussion

EPA decorations facilitate *E. faecalis* virulence by mediating resistance to extracellular stressors and phagocytosis (19). We established an *in vitro* phagocytosis assay using iBMDMs and determined optimum conditions to detect uptake for both wild-type and mutants with an altered cell envelope.

We established that V583 EPA decorations complement OG1RF Δ*epa_var* and vice versa, strongly suggesting that EPA decorations facilitate immune evasion via a conserved mechanism (Fig. 2a-b). The genetic loci encoding EPA decorations in strains OG1RF and V583 are strikingly different. Yet, the expression of both loci in the OG1RF Δ*epa_var* background can inhibit phagocytosis. Structural studies are required to establish if both decorations share motifs sufficient to protect against uptake by macrophages. However, it is tempting to assume that the architecture of EPA, irrespective of its composition, is masking enterococcal cell envelope components from being bound by phagocytic receptors. In future, it would be interesting to expand our EPA cross-complementation study to cover a greater structural diversity of decorations. From this proposed work, it may be possible to define the EPA structural requirements critical for immune evasion or establish that EPA decoration structure is not important so long as a protective barrier is formed.

Our results suggest that the presence of EPA decorations at the cell surface are required to limit bacterial aggregation and thereby minimise the number of bacteria taken up by phagocytes. Given that EPA decorations have a net negative charge (19), we postulate that the decorations reduce aggregation by inhibiting hydrophobic interactions between bacteria. Measurements performed independently by different research groups have consistently shown that the deletion of *epa* genes increases enterococcal surface hydrophobicity (19,30,31). The cell aggregates formed by Δ*epa_var* are more efficiently internalised by macrophages. This conclusion is supported by the fact that GFP-expressing Δ*epa_var* bacteria increase macrophage fluorescence without increasing the percentage of macrophages positive for bacteria (Fig. S4). Dispersion of bacterial aggregates by sonication significantly reduces internalisation. This result is consistent with a previous study showing that the minimization of bacterial cell size is an important factor for the dissemination of *E. faecalis* in the host (13). A similar mechanism has been reported for *Streptococcus pneumoniae*, which also minimises phagocytic uptake by minimising aggregation (32). In contrast, uropathogenic *E. coli* seem to inhibit phagocytosis by morphing into long, filamentous cells whose elongated shape makes phagocytic cup formation less mechanistically favourable (33,34). This illustrates the diversity of mechanisms evolved by bacteria to circumvent phagocytosis.

In the absence of EPA, the enterococcal cell surface can be readily recognised by iBMDMs. Our study indicates that the PAMP(s) responsible for this recognition are not membrane-anchored lipoproteins and therefore remain to be identified. These could be cell wall-anchored proteins, peptidoglycan, rhamnan, or lipoteichoic acids (LTAs). Testing the contribution of some of these components to phagocytic uptake will be challenging. In the presence of LTA synthase (LtaS) inhibitors, *Enterococcus faecium* cells displayed severe growth and morphological defects (35), suggesting that LTAs are essential for this genus. Attempts to delete both enterococcal homologues of LtaS in *E. faecalis* were unsuccessful, (data not shown), further suggesting that LTAs are essential in enterococci. A different approach to modulate the abundance of LTAs may represent an alternative strategy to test.

The scope of this study was limited to non-opsonic phagocytosis. It has been shown elsewhere that an *E. faecalis* V583 EPA decoration mutant is more readily bound by two complement components – mannose-binding lectin and C3b – leading to increased neutrophil-mediated opsonophagocytosis (18,36). Therefore, it would be interesting to investigate the mechanisms (if any) by which EPA decorations in other strains inhibit this process.

The assay described in this study represents a tool to explore the contribution of cell envelope components to innate immune evasion and recognition by phagocytes (Fig. 5). This versatile assay can be used for several purposes: (i) to identify the Pathogen Associated Molecular Patterns (PAMPs) recognized by phagocytes (Fig. 5a), looking for a decreased uptake of mutants built in the OG1RF Δ*epa_var* background; (ii) to explore EPA decorations structure/function (Fig. 5b) and (iii) to test the biological activity of EPA decorations produced by *E. faecalis* isolates (Fig. 5c).

**Fig. 5:**
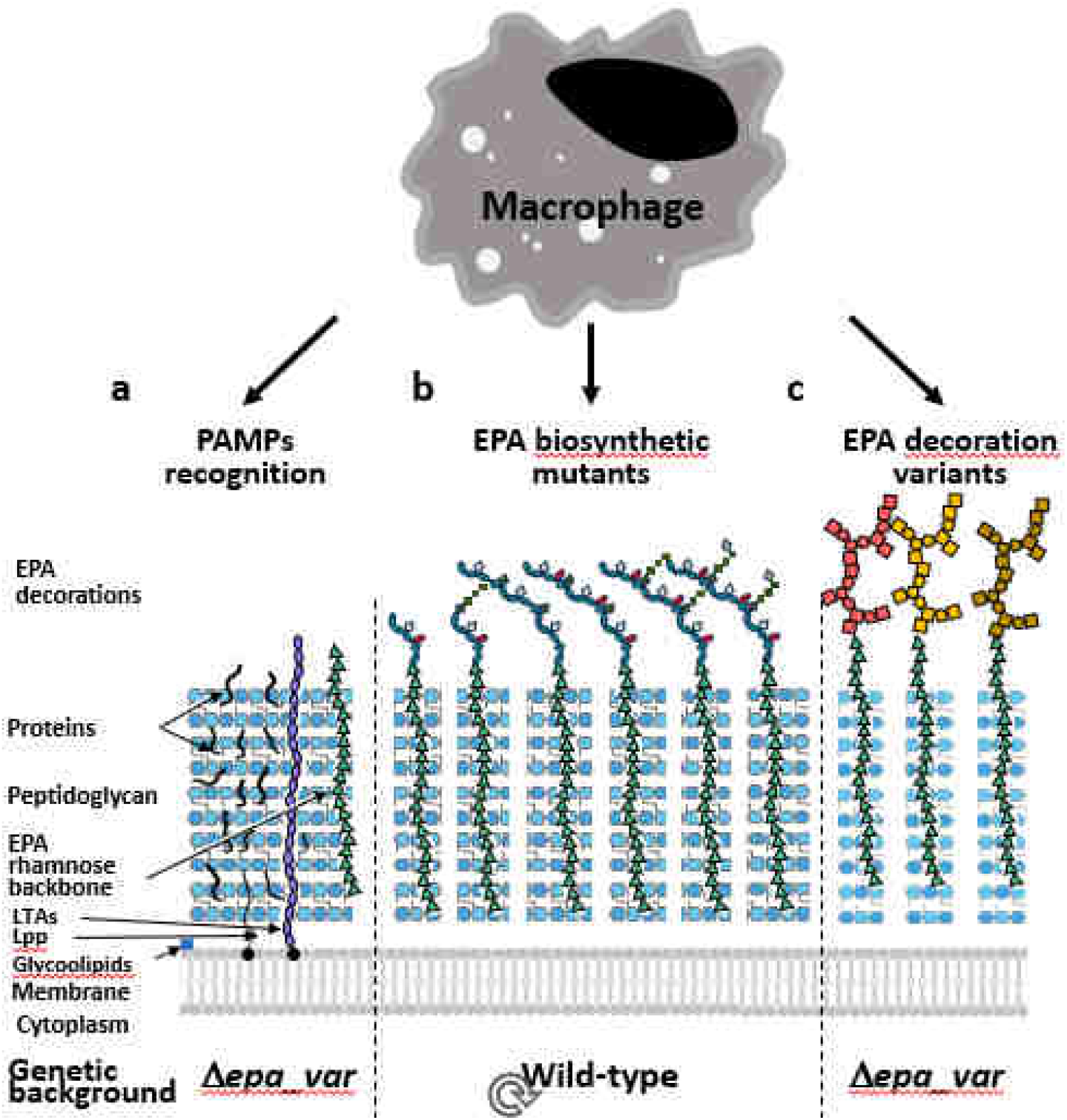
A phagocytosis assay to explore *E. faecalis* interaction with innate immune cells. The assay described in this study can be used to identify the PAMPs recognized by phagocytes (**a**), looking for a decreased uptake of mutants built in the OG1RF Δ*epa_var* background. The analysis of EPA structure/function using NMR and the phagocytosis assay (**b**) will provide insights into the biosynthesis of decorations and the specific contribution of structural determinants to innate immune evasion. The OG1RF Δ*epa_var* can also be used to test the biological activity of EPA decorations produced by *E. faecalis* isolates (**c**).

## Experimental Procedures

### Bacterial strains and growth conditions

All bacterial strains used in this work are listed in Table S1. Unless stated otherwise, *E. faecalis* was cultured by inoculating a single colony into Brain Heart Infusion (BHI) broth and incubating at 37°C without agitation. *E. faecalis* colonies were cultivated on 1.5% (w/v) BHI agar plates at 37°C; these plates were stored at 4°C for up to one month. When appropriate, media/agar was supplemented with antibiotics to maintain selection of plasmids (Table S1). To promote the expression of genes on pTetH2op derivatives, anhydrotetracycline (ATc) was added to a final concentration of 10 ng μl^−1^. *E. coli* work was performed as follows: Unless stated otherwise, a single colony was inoculated into BHI or Luria-Bertani (LB) broth for incubation at 37°C with agitation. Single colonies were cultivated on 1.5% (w/v) BHI agar plates at 37°C. When appropriate, antibiotics were added as described in Table S2.

### Construction of GFP-expressing *E. faecalis*

The plasmid pMV_GFP was electroporated into *E. faecalis* electrocompetent cells. Transformants were selected for on BHI agar + 5 μg/ml Tet plates at 37°C. GFP expression was verified by imaging patched transformants on a Gel DocTM XR+ imager (Alexa488 channel).

### Tissue culture

Immortalised bone marrow-derived macrophages (iBMDMs) from oncogenic mice (21) were cultured in DMEM (Gibco) supplemented with 1% (v/v) foetal bovine serum (FBS, PAN Biotech; low endotoxin, heat inactivated), penicillin (10U/ml)/streptomycin (1mg/ml) (Lonza) and 1% (v/v) sodium pyruvate (Thermo Fisher, 1mM final concentration). Cells were cultured in standard tissue culture flasks or multi-well plates at 37°C in 5% CO_2_, washed in PBS and given fresh media once every 48 hours. Cells were split when >70% confluence had been reached.

### *In vitro* internalisation assay optimisation

1 hour versus 3 hours experiment: on day 1, iBMDMs were checked for >70% confluence. An exact cell count was made using a Countess automated cell counter (Invitrogen) as per the manufacturer’s instructions. iBMDMs were diluted to 4 x 10^5^ live cells per ml in fresh media. To set up one technical replicate, a 5 ml (2 x 10^6^ cells) was transferred to a 25 ml tissue culture flask. Flasks were incubated as normal. In addition, one *E. faecalis* OG1RF pMV_GFP overnight culture was set up in 10 ml BHI + 5 μg/ml Tet. On day 2, iBMDMs were washed once with PBS and given 1 ml fresh DMEM (serum- and antibiotic-free). Bacteria were harvested (5 min at 4000 x g) and resuspended in PBS. Optical density of bacterial suspensions was normalised to OD600 = 1 (1 x 10^9^ CFU/ml). Bacterial suspension was 10x diluted in DMEM (serum- and antibiotic-free), giving 1 x 10^8^ CFU/ml. 1 ml this suspension was added to each iBMDM flask (MOI = 50), then flasks were incubated for 1 hour or 3 hours at 37°C, 5% CO2. For the 3 hours on ice control, flasks were incubated on ice for 5 min prior to addition of bacteria. Post-incubation, iBMDMs were washed three times with PBS and treated with 5 ml DMEM + 250 μg/ml gentamycin + 20 μg/ml vancomycin for 1 hour at 37°C, 5% CO2. Then, iBMDMs were washed twice with PBS, resuspended in 5 ml PBS using a cell scraper, and transferred to 15 ml Falcon tubes. iBMDMs were pelleted (5 min at 4000 x g) and resuspended in 1 ml PBS + 4% (m/v) paraformaldehyde for 10 min fixing at room temperature. iBMDMs were re-pelleted, washed once with PBS, resuspended in 500 μl filtered-sterilised PBS, and stored at 4°C in darkness until day 3.

MOI dose response experiment: the method used was mostly the same as described above, but with the changes outlined here: On day 1 confluent iBMDMs were diluted to 1 x 10^5^ live cells/ml. Two ml (2 x 10^5^ iBMDMs) were transferred to each well a six-well plate. On day 2, an *E. faecalis* suspension was prepared as above, and increasing volumes were added to iBMDMs to give MOI = 1, 5, 10, 20 or 100. The volume of DMEM added (serum- and antibiotic-free) added to each well was adjusted so total volume = 2 ml. Incubation time = 1 hour (37°C, 5% CO2). In total, three six-well plates were used to perform three technical replicates per MOI plus bacteria-free control wells. After incubation, the method used was the same as described above, except that (i) the volume of DMEM + gentamycin + vancomycin was 2 ml per well, (ii) the volume of PBS used for washing/resuspending was 1 ml, and (iii) the volume of 4% PFA used for fixing was 500 μl.

### *In vitro* internalisation assays to compare uptake of different *E. faecalis* strains

On day 1, iBMDMs were checked for >70% confluence and counted. iBMDMs were diluted to 2.5 x 10^5^ live cells per ml in fresh media; 5 x 10^5^ cells were aliquoted per well. *E. faecalis* overnight cultures were set up as standard. On day 2, fresh *E. faecalis* cultures were started (100 μl overnight into 10 ml fresh media) and grown at 37°C until OD_600_ ≈ 0.3. Cultures were pelleted (5 min at 4000 x g) and resuspended in an equal volume of DMEM (serum- and antibiotic-free). iBMDMs, after being washed and given fresh media as before, were given 2.5 x 10^6^ CFU bacteria per well (MOI = 5). Three wells were allocated per bacterial strain in each experiment. After 1 hour at 37°C in 5% CO_2_, iBMDMs were washed and treated with antibiotics as before. Then, cells were washed twice with PBS and detached by treating with 1 ml Accutase^TM^ (Merck) for 30 min at 37°C in 5% CO_2_. Detached iBMDMs were pelleted (5 min at 7,000 x g), fixed, resuspended in 200 μl filtered PBS, and stored at 4°C in darkness.

### Sonication of *E. faecalis* Δ*epa_var* cells

A 5 ml aliquot of each bacterial suspension was treated with 20 cycles of sonication (5 seconds at 20% amplitude) using a Fisherbrand^TM^ 505 sonicator (Fisher).

### Flow cytometry analysis of iBMDMs

200 μl iBMDM samples were vortexed gently and transferred to a 96-well plate. Data acquisition was performed using a Guava easyCyte HT flow cytometer (Luminex). Data analysis was carried out using guavaSoft version 3.1.1; gating strategy is shown in Fig. S1.

### Fluorescence microscopy of iBMDMs

*In vitro* phagocytosis assay was performed exactly as above. 100 μl iBMDM samples were transferred to a 24-well plate. Each sample was diluted by adding 1 ml PBS. iBMDMs images were captured with Elements software (Nikon) using an Andor Neo camera on Nikon Ti microscope with differential interference contrast (DIC) and GFP epifluorescence. In (Fiji is just) ImageJ version 2.9.0/1.5t, iBMDMs that had overlapping GFP signals were identified, and the fluorescence was quantified as mean gray value (MGV). MGV measurements were normalised by subtracting the average MGV of the background of the image. For each group, >90 macrophages were measured.

### *E. faecalis* growth curves

*E. faecalis* overnight cultures (three per strain) were set up as normal. The next morning, each overnight culture was serially diluted in a 96 well plate. Each dilution step meant transferring 20 μl culture to 180 μl fresh BHI broth (i.e., a 10-fold dilution). After the final dilution, each culture had been diluted by 1 x 10^3^. The plate’s lid was replaced only after it had been treated with a solution of 0.05% (v/v) Triton X-100 + 20% v/v ethanol to prevent condensation. The plate was loaded into a Sunrise^TM^ microplate reader (Tecan), and growth was allowed to proceed for 24 hours at 37°C. Optical density (OD) measurements were taken every 5 mins (wavelength = 595 nm). Cultures were agitated for 5 s at normal power before each measurement. Once the run had been completed, each curve was plotted as OD_595_ (y axis, logarithmic) versus time in minutes (x axis, linear).

### CFU/ml determination of exponential *E. faecalis* cultures

*E. faecalis* cultures were set up by using 100 μl overnight culture to inoculate 10 ml fresh BHI broth. Cultures were incubated at 37°C without agitation until OD_600_ ≈ 0.3. Ten-fold serial dilutions were performed in PBS until the cultures had been diluted by 1 x 10^7^. 100 μl of each final dilution was plated onto a standard BHI agar plate and incubated overnight at 37°C. The next morning, each plate was placed under a Scan4000 automated colony counter (Interscience). By calculating backwards from the CFU counts, CFU/ml values of the undiluted cultures were determined and normalised to OD_600_ = 0.3. A mean CFU/ml value was determined for each strain from at least three independent cultures.

### Fluorescence microscopy of *E. faecalis*

*E. faecalis* was grown until OD_600_ ≈ 0.3. One ml of each culture was pelleted (6,000 x g, 1 min), resuspended in 1 ml leftover culture, and stained with 5 μl of 50mM HADA (10 min on a rotary shaker at 37°C in complete darkness). Bacteria (kept wrapped in foil to prevent photobleaching) were pelleted as before, washed twice with PBS, and resuspended in 300 µl of PBS. Next, bacteria were supplemented with 5 μl of AlexaFluor^TM^ 555 NHS ester (Molecular Probes) at a concentration of 1 mg/ml and left to be stained for 7 min at room temperature. As a fixing step, bacteria were pelleted, resuspended in 750 μl 4% (m/v) paraformaldehyde in PBS, and left for 30 min at room temperature. After fixing, cells were washed twice in PBS and resuspended in 20 µ l of MilliQ water. Five μl were mounted onto a PolyPrep slide using SlowFade^TM^ Gold (Thermo Fisher) and a standard 13 mm coverslip. Images were captured on a Nikon DualCam system (Eclipse Ti inverted research microscope). Wavelengths and filters (Table S3) were applied as appropriate for each image. Contrast and brightness adjustments were made in ImageJ.

### Phase contrast microscopy of *E. faecalis*

*E. faecalis* strains were grown to OD_600_ ≈ 0.3, then a 1 ml aliquot of each culture was pelleted (6,000 x g, 1 min). Bacteria were fixed in 750 μl 4% (m/v) paraformaldehyde as described in the previous section. Fixed bacteria were washed twice in PBS, resuspended in 20 µl of MilliQ water, and mounted as described previously. Images were captured on a Nikon DualCam system (Elipse Ti inverted research microscope). Cell length and width measurements were made using the ObjectJ plugin in ImageJ.

### Flow cytometry analysis of *E. faecalis*

*E. faecalis* strains were grown to OD_600_ ≈ 0.3, pelleted (4,000 x g, 5 min) and resuspended in PBS at an OD_600_ of 0.4. Bacterial suspensions were treated with 10, 20, and 30 pulses using the Fisherbrand^TM^ 505 sonicator. After every 10 pulses, 200 μl of the cell suspension were taken out and serially diluted (10-fold dilutions until diluted by 1 x 10^4^). To measure CFU/ml, two 100 μl aliquots of each final dilution were plated and CFU were counted using the Scan4000 colony counter (Interscience). To measure the forward scatter (FSC), 10 x dilutions of bacterial suspensions were passed through the Guava easyCyte HT, followed by data processing and analysis as described in Fig. S8. To provide a negative control, we also analysed some bacterial suspension that was set aside and not sonicated.

### Construction of pG_lgt for allelic replacement

Plasmids and oligos used in this study are listed in Table S1. Two homology regions flanking the *lgt* open reading frame were amplified from OG1RF genomic DNA via PCR. The 5’ arm (∼0.75 kb) was amplified using the primers SM_0194 (sense) and SM_0195 (antisense), whereas the 3’ arm (∼0.75 kb) was amplified using SM_0196 (sense) and SM_0197 (antisense). Once purified, the two PCR products were mixed (equimolar amount of each) and fused into a single product (∼1.5 kb) via splice overlap extension PCR (Ho et al., 1989) using primers SM_0194 and SM_0197. The resulting fragment was cut by *Xho*I and *Not*I and cloned into pGhost9 vector cut with the same enzymes (37). Candidate pGhost derivatives were screened by PCR using primers SM_0171 and SM_0172. A positive clone containing the fused H1-H2 insert was checked by sanger sequencing and the corresponding plasmid was named pG_lgt.

### Construction of *E. faecalis* Δ*lgt* mutants

*E. faecalis* mutants were built by allelic exchange as previously described (38). Purified pG_lgt plasmid was electroporated into *E. faecalis* OG1RF wild type and OG1RF Δ*epa_var*. Transformants were selected on BHI agar + 30 μg/ml erythromycin plates at 28°C (a plasmid replication-permissive temperature). Transformants were then streaked onto BHI agar 30 μg/ml erythromycin without antibiotic at 42°C (a non-replication-permissive temperature) to select plasmid single crossover recombination events. Colonies from these plates were used to inoculate BHI broth cultures and passaged repeated at 28°C without antibiotic. To find double crossover recombination events, single colonies were re-isolated and screened via PCR using the primers SM_0210 and SM_0211 (Table S1). Double crossovers – corresponding to Δ*lgt* mutants – were identified in both backgrounds and validated by purifying their genomic DNA sequencing the *lgt* locus.

### Complementation of *E. faecalis* Δ*lgt* mutants

The complete *lgt* gene was PCR amplified using primers SM_0401 and SM_0402 and cloned into the pTetH vector using NcoI and BamHI. Candidates were screened by using primers SM_0100 and SM_0101. A positive clone containing the *lgt* insert was checked by sanger sequencing and the corresponding plasmid was named pTetH_lgt. *lgt* expression was induced by adding 10 ng/ml anhydrotetracycline.

### Preparation of protein extracts from *E. faecalis* culture supernatants

*E. faecalis* was grown to OD_600_ ≈ 0.3 as normal. Proteins contained in 1.8 ml of culture were precipitated by addig 200 μl 100% (w/v) trichloroacetic acid (TCA). After 15 min on ice, the samples were spun (25,000 x g, 10 min at 4°C). Proteins were washed in μl acetone, centrifuged (25,000 x g, 5 min at 4°C) and left to dry. Pellets were resuspended in 95 μl PBS + 5 μl Tris base and stored at −80°C.

### SDS-PAGE

SDS-PAGEwas performed as previously described (39) Protein extracts were mixed with 5x loading dye (250 mM Tris-HCl (pH 6.8), 10% (w/v) SDS, 0.5% (w/v) bromophenol blue, 50% (v/v) glycerol, 0.5 M dithiothreitol). Gels were stained with a Coomassie solution (0.25% (w/v) Coomassie blue R-250, 50% (v/v) methanol, 10% (v/v) glacial acetic acid) for 1 hour at room temperature with gentle rocking and destained in 5% (v/v) methanol, 10% (v/v) glacial acetic acid).

### Statistical analysis

GraphPad Prism version 10.1.2 was utilised for statistical analysis. Unless stated otherwise, all error bars on graphs represent mean ± SD. Each set of iBMDM flow cytometry data was analysed using a one-way ANOVA with Welch’s correction, followed by Dunnett’s multiple comparisons test. *E. faecalis* doubling times, CFU/ml at early exponential phase, *E. faecalis* cell length/width, and *E. faecalis* FSC between strains were also analysed in this manner. Since there were only two groups in the 37°C versus 4°C phagocytosis assay, an unpaired, two-tailed *t-*test with Welch’s correction was used here. iBMDM microscopy data was analysed using a Kruskal-Wallis test followed by Dunn’s multiple comparisons test. For the experiment comparing *E. faecalis* strain versus FSC versus number of pulses, a two-way ANOVA was performed, followed by Tukey’s multiple comparisons test.

## Acknowledgements

We thank Pascale Serror for sharing the OG1RF Δ*epa_var* strain. JSN was supported by a studentship from the DiMeN Doctoral Training Programme (Medical Research Council grant MR/N013840/1). Jessica Davis was funded by the White Rose Doctoral Training Programme (BBSRC grant BB/ M011151/1).

## Author contributions

**Conceptualization:** JSN, SM, EKT; **Data Curation:** JSN, SM, SAJ, EKT; **Formal Analysis:** JSN; **Funding Acquisition:** SM; **Investigation:** JSN, JLD, BS, CEM; **Methodology:** JSN, PEE, SAJ, ETK; **Project Administration:** PEE, ETK, SAJ, SM; **Resources:** SAJ, CEM, ETK; **Supervision:** SAJ, PEE, EKT, SM; **Validation:** JSN, ETK, SM; **Visualization:** JSN, JLD, SM; **Writing – Original Draft Preparation:** JSN, SM; **Writing – Review & Editing:** JSN, JLD, CEM, PEE, EKT, SAJ, SM

## Supplementary materials

Supplementary Tables: S1, S2 and S3

Supplementary Figures: S1, S2, S3, S4, S5, S6, S7 and S8

## Data availability

Raw data and materials described in this study are available upon request.

---

**Fig. S1:**
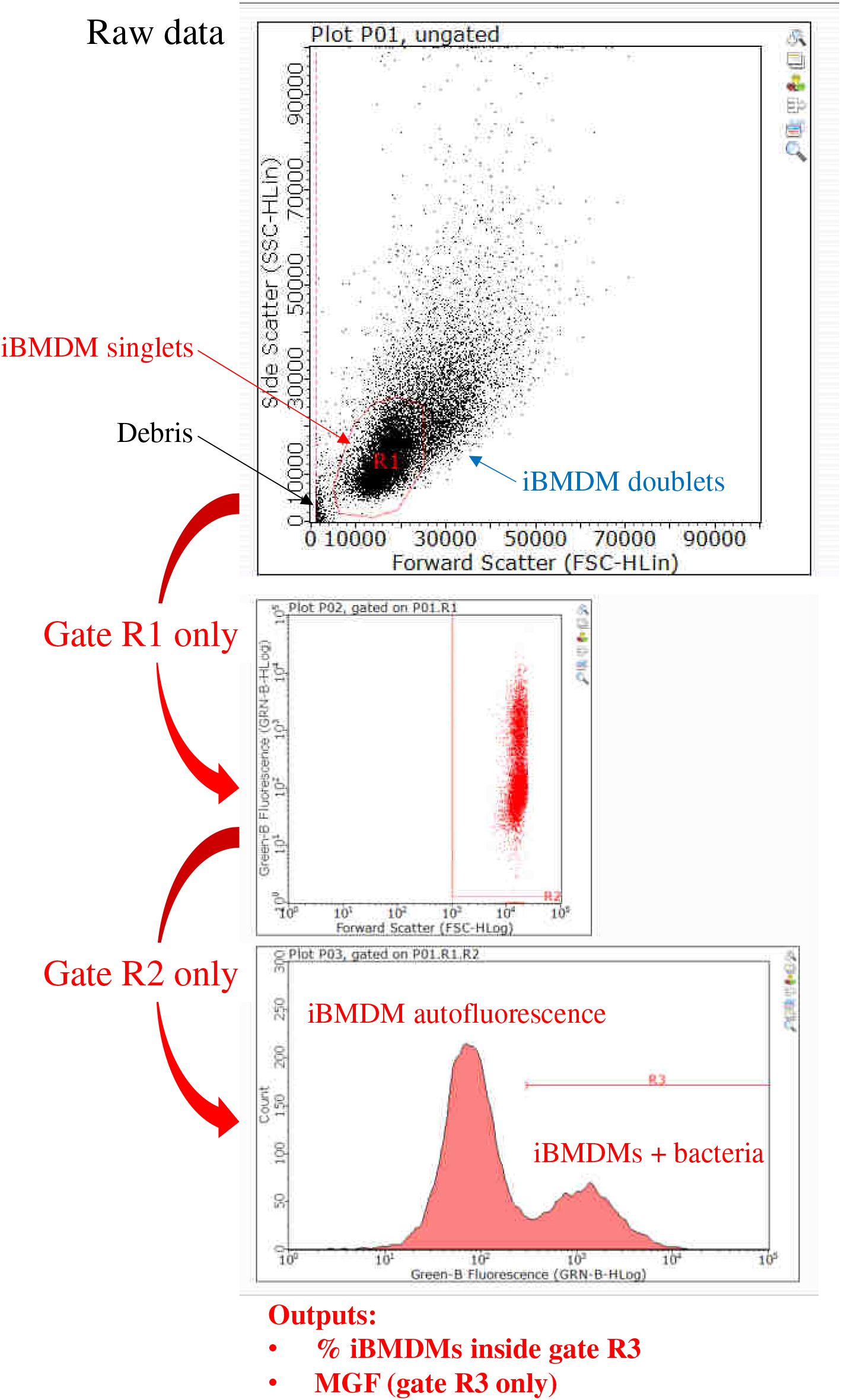
Gating strategy for flow cytometry analysis of iBMDMs using GuavaSoft 3.1.1. Debris and cell clumps were excluded from gate R1. Gate R1 data was re-plotted as FSC log (x axis) versus green fluorescence log (y axis). Gate R2 excluded more debris. Gate R2 data was plotted as a histogram (green fluorescence log (x axis) versus count (y axis)). The left peak (peak green fluorescence ≈ 7 x 10^1^) corresponds to autofluorescence of empty iBMDMs, whereas the right peak corresponds to iBMDMs with internalised GFP-labelled bacteria. Gate R3 (green fluorescence > 3 x 10^2^) was drawn to select only the right peak. The percentage of iBMDMs containing bacteria was calculated using (no. iBMDMs in gate R3/no. iBMDMs in gate R2) x 100. The median green fluorescence (MGF) of gate R3 data was calculated by GuavaSoft 3.1.1.

**Fig. S2:**
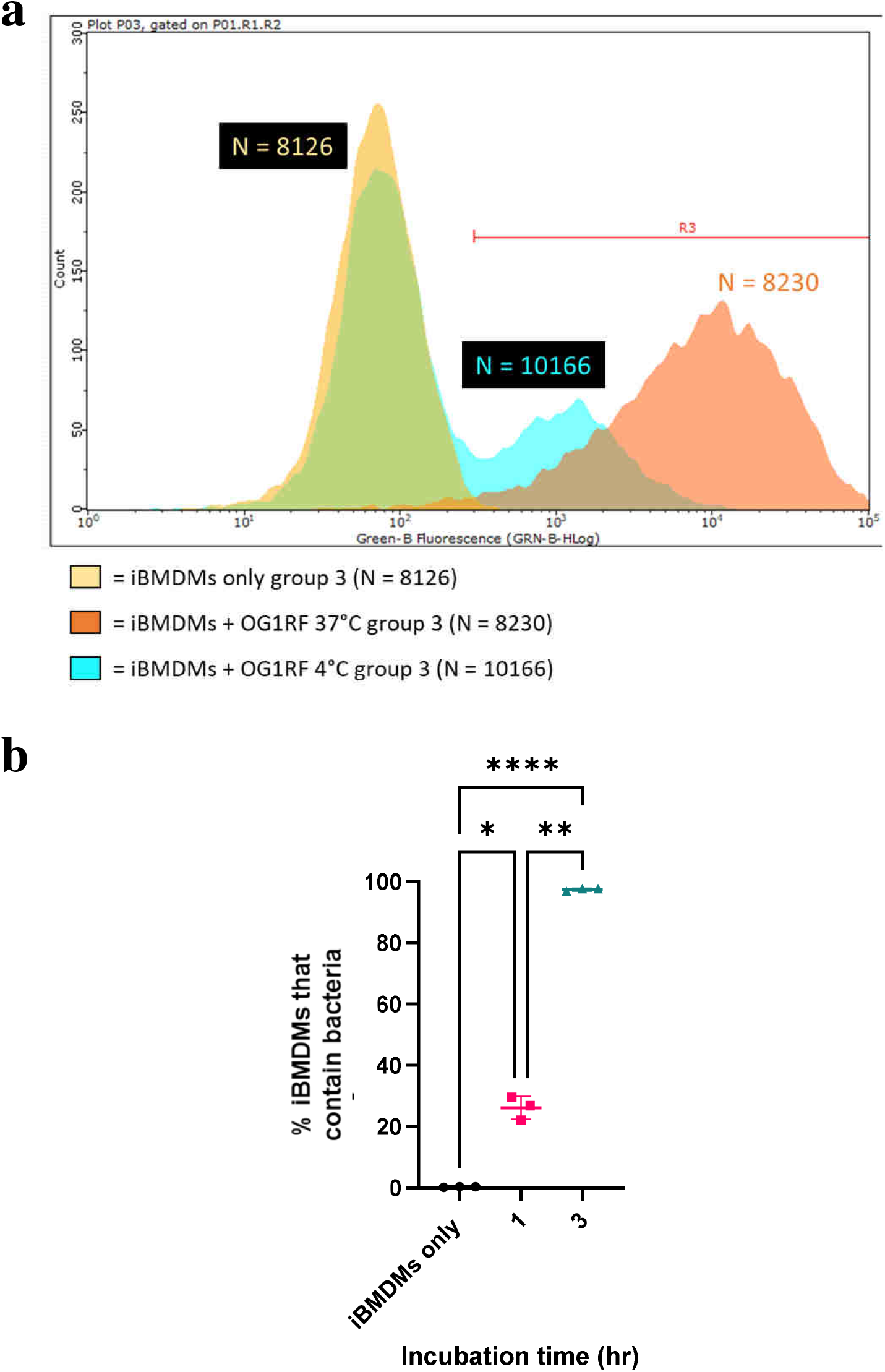
Impact of incubation time and multiplicity of Infection (MOI) on *E. faecalis* uptake by iBMDMs. **(a)** Histograms plotting green fluorescence of iBMDMs following incubation without treatment (yellow) or with GFP-labelled OG1RF for 1 hr (blue) or 3 hr (orange) at 37°C. The position of gate R3 (which contains bacteria-containing macrophages) is indicated. Each plot represents one of three independent replicates performed for each treatment. In this figure, N = total number of iBMDMs per group. **(b)** Percentage of iBMDMs that contain *E. faecalis* after 1 hour versus 3 hours incubation at 37 °C. A one-way ANOVA with Brown-Forsythe and Welch’s correction followed by Dunnett’s multiple comparisons test was performed to assess significance. *P*-values: iBMDMs only versus 1 hour, *P* = 0.0146; iBMDMs only versus 3 hours, *P* < 0.0001; 1 hour versus 3 hours, *P* = 0.002. Error bars represent mean ± standard deviation (SD). *P*-value descriptors: *, *P* < 0.05; **, *P* < 0.01; ****, *P* < 0.0001.

**Fig. S3:**
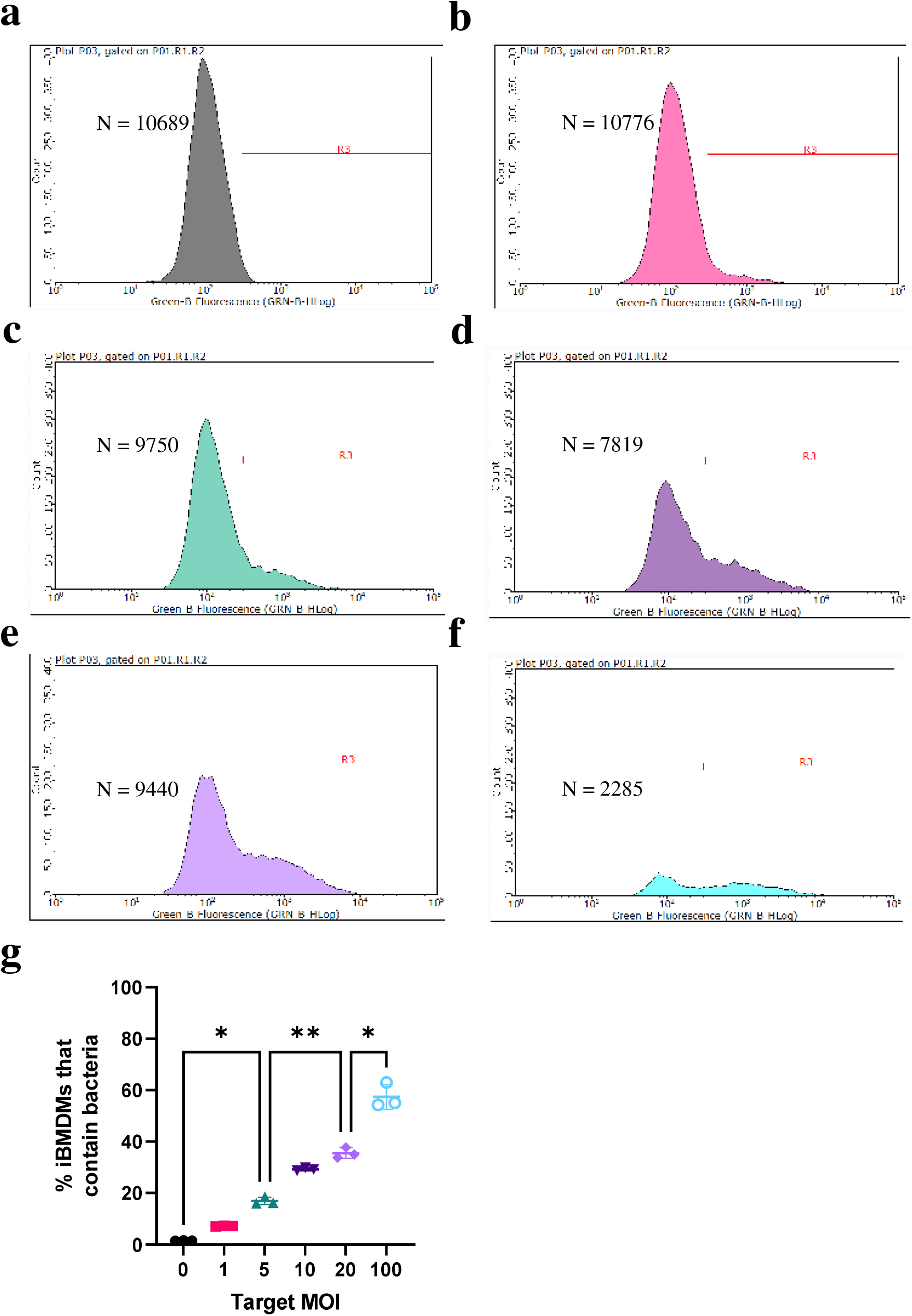
Impact of multiplicity of Infection (MOI) on *E. faecalis* uptake by iBMDMs – histograms and proportions. **(a-f)** Histograms plotting green fluorescence of iBMDMs following incubation without treatment **(a)** or with GFP-labelled OG1RF at MOI = 1 **(b)**, 5 **(c)**, 10 **(d)**, 20 **(e)**, or 100 **(f)**. On each plot, the position of gate R3 (which contains bacteria-containing macrophages) is indicated. Each plot represents one of three independent replicates performed for each MOI. In this figure, N = total number of iBMDMs per plot. (**g**) Percentage of iBMDMs that contain *E. faecalis* according to bacterial dose (1 hour incubation). Statistical analysis was performed via a one-way ANOVA with Brown-Forsythe and Welch’s correction followed by Dunnett’s multiple comparisons test. *P*-values: MOI = 0 versus MOI = 5, *P* = 0.0126; MOI = 5 versus MOI = 20, *P* = 0.0016; MOI = 20 versus MOI = 100, *P* = 0.0329. Error bars represent mean ± standard deviation (SD). *P*-value descriptors: *, *P* < 0.05; **, *P* < 0.01.

**Fig. S4:**
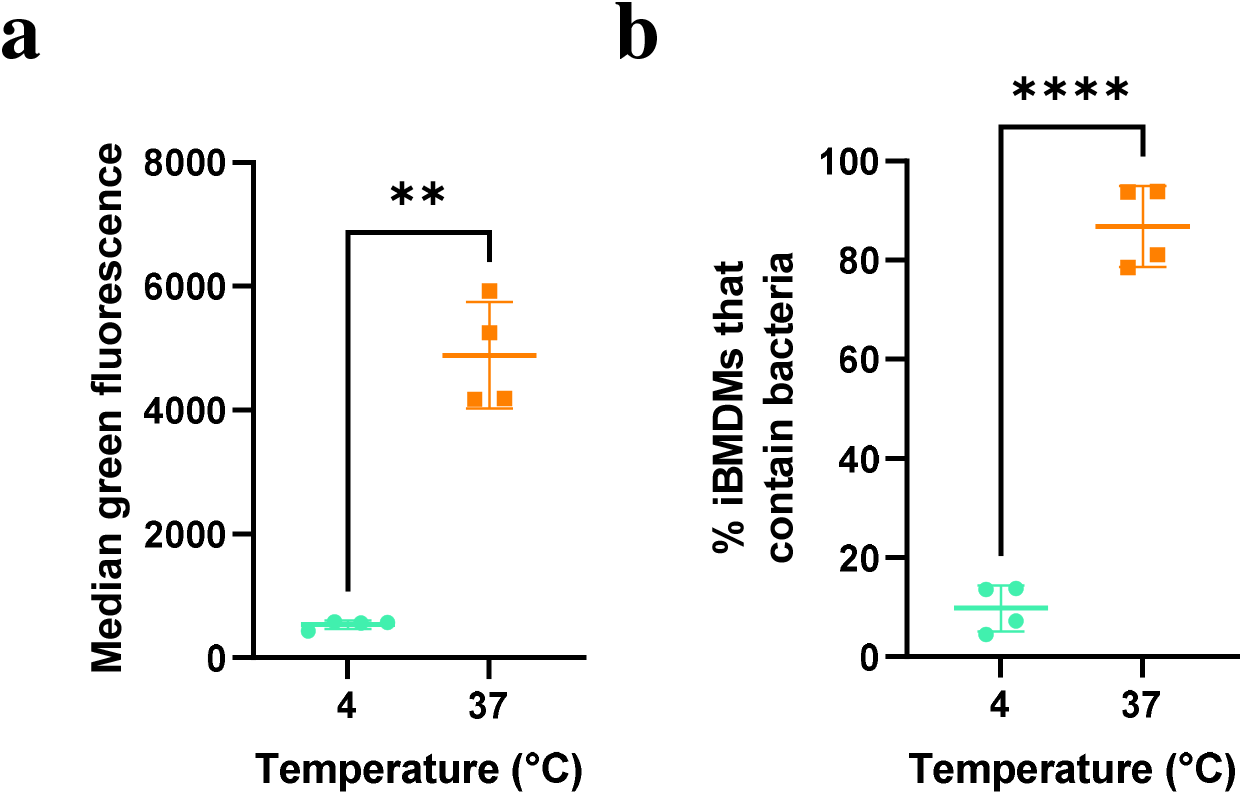
Impact of temperature on *E. faecalis* uptake by iBMDMs. (**a**) *E. faecalis*-positive iBMDMs were significantly more fluorescent at 37 °C as compared to 4 °C. Statistical analysis was performed via an unpaired *t*-test with Welch’s correction (*P* = 0.0019; n = 4 technical replicates). (**b**) The percentage of iBMDMs that contained bacteria was found to be significantly higher at 37 °C as compared to 4 °C. An unpaired *t*-test was performed with Welch’s correction (*P* < 0.0001; n = 4 technical replicates).

**Fig. S5:**
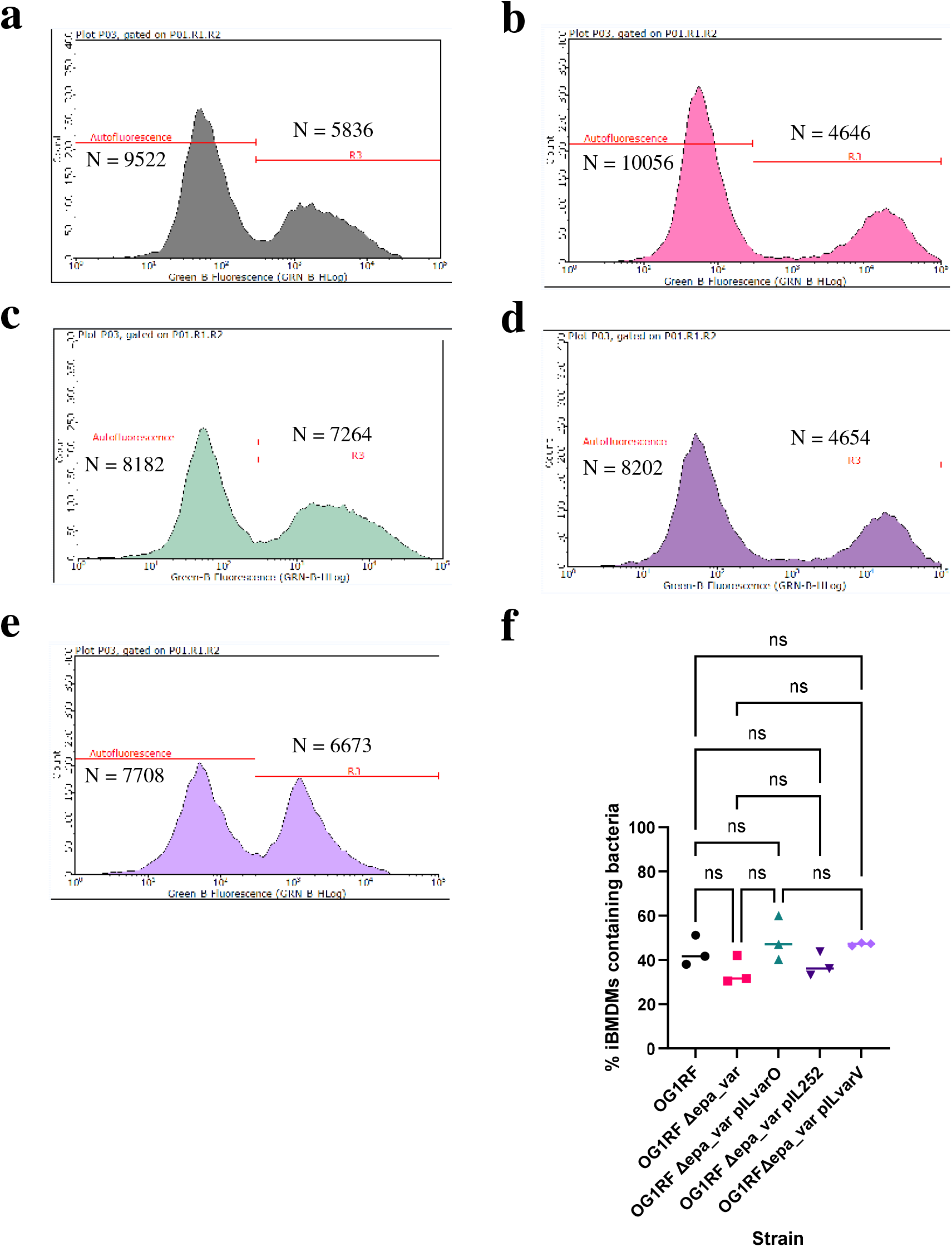
Phagocytosis of *E. faecalis* OG1RF derivatives – histograms and proportions. **(a-e)** Histograms plotting green fluorescence of iBMDMs following incubation with GFP-labelled OG1RF **(a)**, the Δ*epa_var* derivative **(b)**, Δ*epa_var* pILvarO **(c)**, Δ*epa_var* pIL252 **(d**), or Δ*epa_var* pILvarV **(e)**. On each plot, the separation between bacteria-free (Autofluorescence) and bacteria-positive (gate R3) macrophages is indicated. Each plot represents one of three independent replicates performed for each treatment in this experiment. In this figure, N = number of iBMDMs within each gate. **(f)** Percentage of iBMDMs that did contain bacteria. Each value is the mean of three replicates per treatment. Statistical analysis was performed by via one-way ANOVA with Brown-Forsythe and Welch’s correction followed by Dunnett’s multiple comparisons test. *P*-values: OG1RF versus Δ*epa_var*, *P* = 0.664; OG1RF versus pILvarO, *P* = 0.985; OG1RF versus pIL252, *P* = 0.890; OG1RF versus pILvarV, *P* = 0.952; Δ*epa_var* versus pILvarO, *P* = 0.491; Δ*epa_var* versus pIL252, *P* = 0.997; Δ*epa_var* versus pILvarV, *P* = 0.270; pILvarO versus pILvarV, *P* > 0.999. *P*-value descriptors: ns, not significant.

**Fig. S6:**
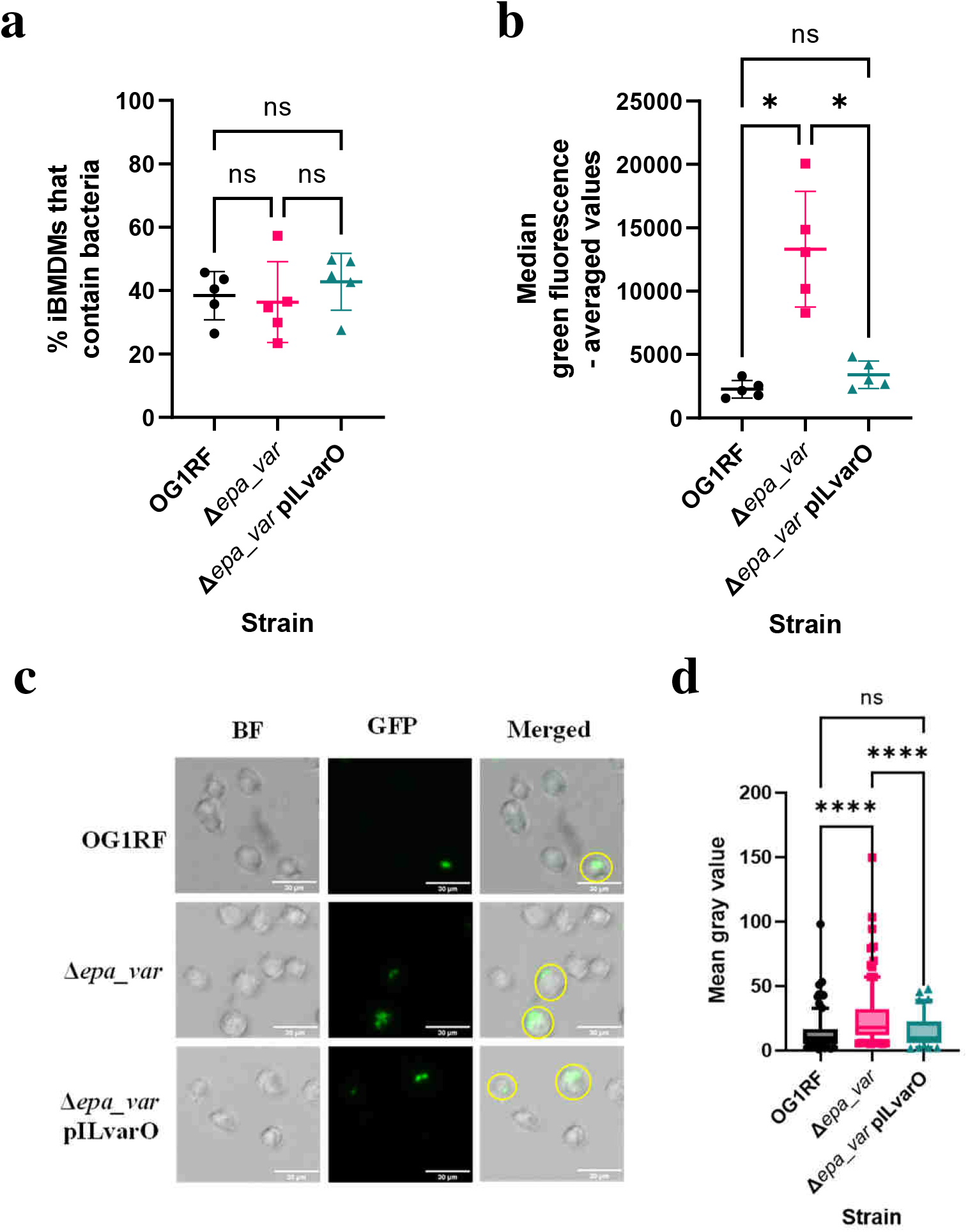
Internalisation of *E. faecalis* OG1RF Δ*epa_var* which lacks EPA decorations. (**a**) Percentage iBMDMs positive for internalised bacteria. *P*-values: OG1RF versus Δ*epa_var*, *P* = 0.985; OG1RF versus pILvarO, *P* = 0.794; Δ*epa_var* versus pILvarO, *P* = 0.743. Statistical analysis was performed by doing a one-way ANOVA with Brown-Forsythe and Welch’s correction followed by Dunnett’s multiple comparisons test (n = 5 biological replicates per group). (**b**) Green fluorescence intensity of iBMDMs that contained bacteria. *P*-values: OG1RF versus Δ*epa_var*, *P* = 0.0150; OG1RF versus pILvarO, *P* = 0.220; Δ*epa_var* versus pILvarO, *P* = 0.0232. Again, statistical analysis was performed via a one-way ANOVA with Brown-Forsythe and Welch’s correction followed by Dunnett’s multiple comparisons test (n = 5 biological replicates per group). (**c**) Confocal microscopy of iBMDMs following incubation with GFP-labelled *E. faecalis* – representative images. Yellow circles indicate macrophages analysed in (**d**). BF, brightfield; GFP, GFP channel. (d) Pixel intensity of macrophages with internalised bacteria, measured as mean grey value using ImageJ. All values were normalised by subtracting the mean grey value of the background. Box plots represent medians flanked by upper and lower quartiles (25^th^ and 75^th^ percentiles, respectively), while whiskers represent 5^th^ and 95^th^ percentiles. Statistical analysis was performed by doing a Kruskal-Wallis test followed by Dunn’s multiple comparisons test. *P*-values: OG1RF versus Δ*epa_var*, *P* < 0.0001; OG1RF versus pILvarO, *P* > 0.999; Δ*epa_var* versus pILvarO, *P* < 0.0001. Sample sizes: OG1RF, n = 203; Δ*epa_var*, n = 215; Δ*epa_var* pILvarO, n = 98. Key to *P*-values: ns, not significant; *, *P* < 0.05; ****, *P* < 0.0001.

**Fig. S7:**
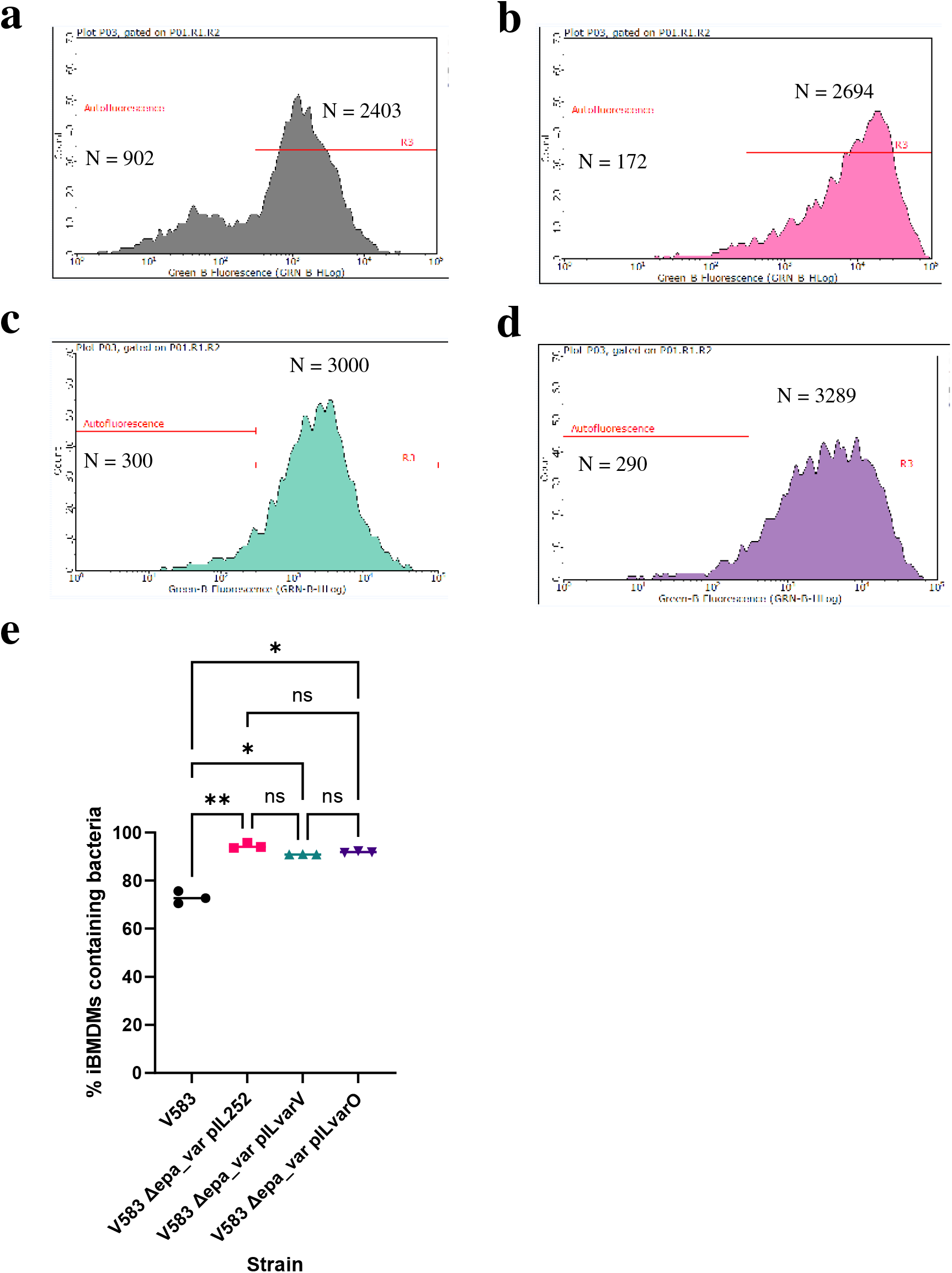
Phagocytosis of *E. faecalis* V583 derivatives – histograms and proportions. **(a-d)** Histograms plotting green fluorescence of iBMDMs following incubation with GFP-labelled V583 **(a)**, or the Δ*epa_var* derivative with pIL252 **(b)**, pILvarV **(c)**, or pILvarO **(d**). On each plot, the separation between bacteria-free (Autofluorescence) and bacteria-positive (gate R3) macrophages is indicated. Each plot represents one of three independent replicates performed for each treatment in this experiment. In this figure, N = number of iBMDMs within each gate. **(e)** Percentage of iBMDMs that did contain bacteria. Each value is the mean of three replicates per treatment. Statistical analysis was performed by via one-way ANOVA with Brown-Forsythe and Welch’s correction followed by Dunnett’s multiple comparisons test. *P*-values: V583 versus pIL252, *P* = 0.0035; V583 versus pILvarV, *P* = 0.0201; V583 versus pILvarO, *P* = 0.0184; pIL252 versus pILvarV, *P* = 0.103; pIL252 versus pILvarO, *P* = 0.201; pILvarV versus pILvarO, *P* = 0.117. *P*-value descriptors: ns, not significant; *, P < 0.05; **, P < 0.01.

**Fig. S8:**
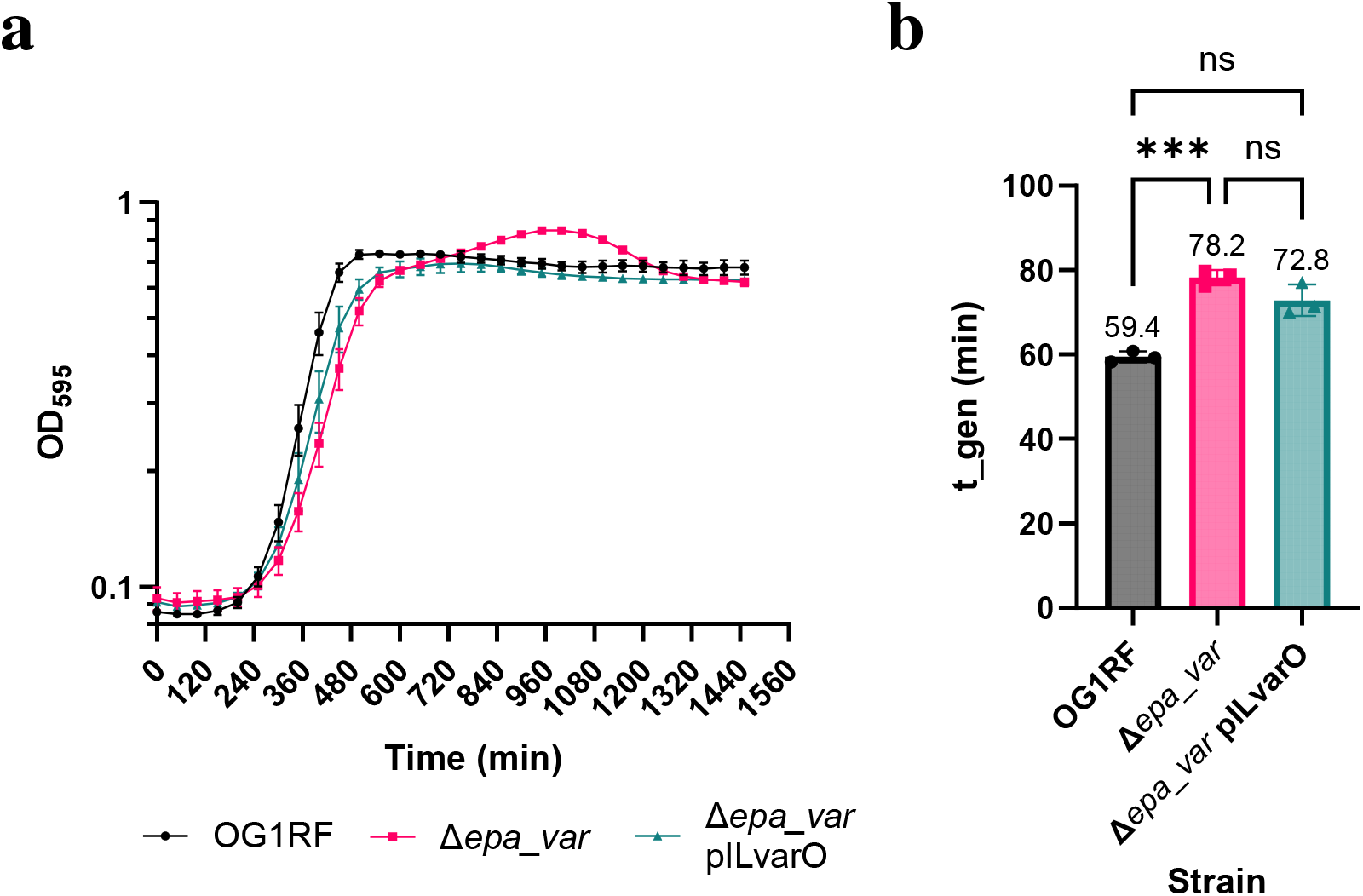
Impact of the Δ*epa_var* mutation on *E. faecalis* OG1RF growth rate. (**a**) Growth profiles of *E. faecalis* OG1RF, Δ*epa_var* and complemented Δ*epa_var* in BHI broth at 37 °C. Each data point represents the mean of three biological replicates ± SD. (**b**) Generation times (t_gen) in min. Three biological replicates per strain were performed. Mean t_gen values were compared via one-way ANOVA with Brown-Forsythe and Welch’s correction, followed by Dunnett’s multiple comparisons test. *P*-values: OG1RF versus Δ*epa_var*, *P* = 0.0003; OG1RF versus pILvarO, *P* = 0.0568; Δ*epa_var* versus pILvarO, *P* = 0.240. Key to *P*-values: ns, not significant; ***, *P* < 0.001.

**Fig. S9:**
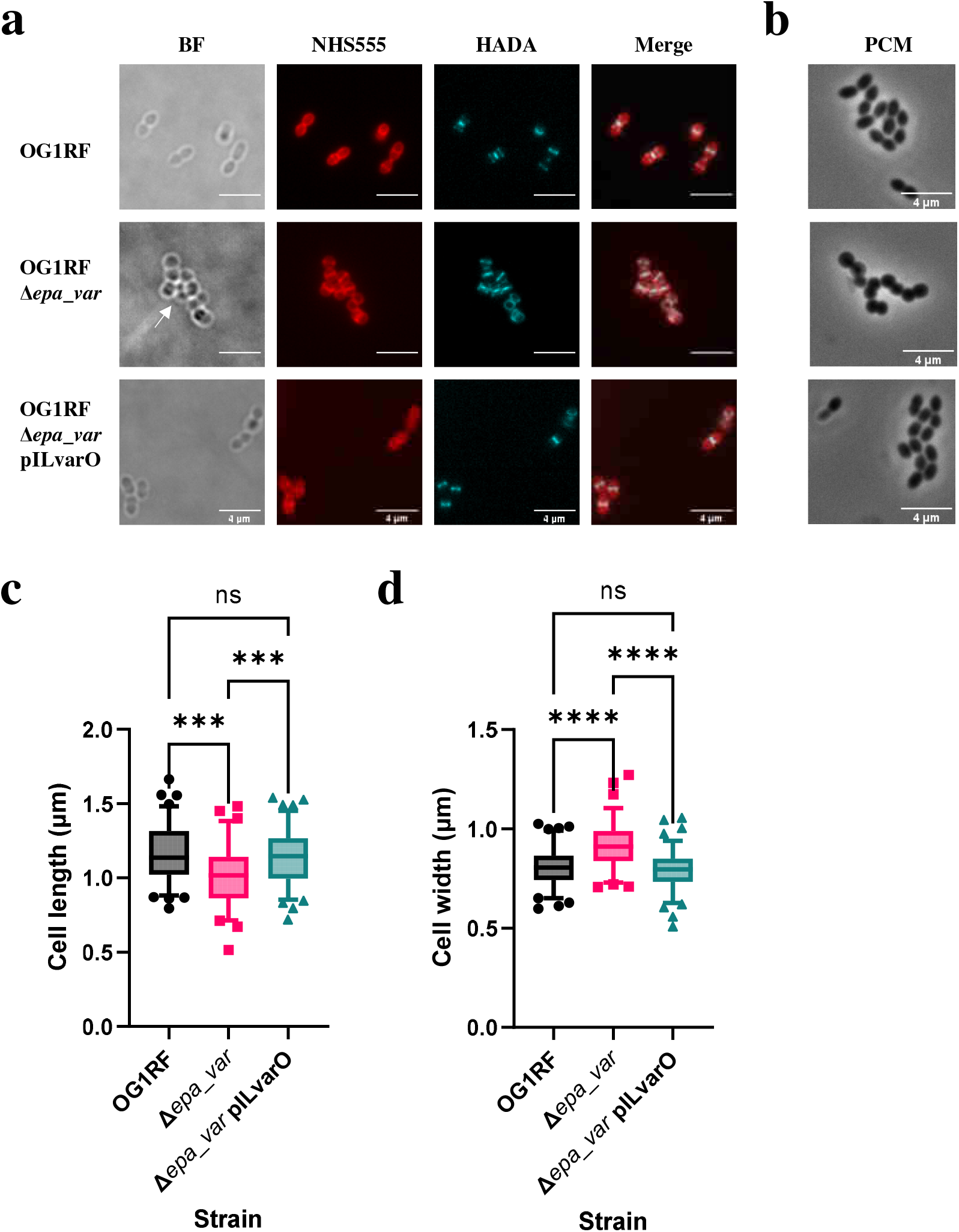
Microscopic analysis of *E. faecalis* OG1RF Δ*epa_var* shows an altered morphology of *epa_var*. (**a**) Fluorescence microscopy of exponential-phase *E. faecalis* bacteria labelled with NHS ester 555 and HADA. White arrows indicate bacterial cell aggregates. All images were taken at 100 x magnification. Scale bar = 4 μm. BF, brightfield. (**b**) Phase contrast microscopy of exponential-phase *E. faecalis*. OG1RF = upper panel; Δ*epa_var* = middle panel; Δ*epa_var* pILvarO = lower panel. Same magnification and scale bar as used in (**a**). (**c**) Comparison of bacterial cell length. Box plots show medians flanked by lower and upper quartiles; whiskers show 5th and 95th percentiles. *P*-values: OG1RF versus Δ*epa_var*, *P* = 0.0002; OG1RF versus pILvarO, *P* > 0.999; Δ*epa_var* versus pILvarO, *P* = 0.0002. Sample sizes: n = 86 (OG1RF); n = 74 (Δ*epa_var*); n = 96 (Δ*epa_var* pILvarO). (**d**) Comparison of bacterial cell width. The same samples were analysed here as (**c**). Box plots show medians flanked by lower and upper quartiles; whiskers show 5th and 95th percentiles. *P*-values: OG1RF versus Δ*epa_var*, *P* < 0.0001; OG1RF versus pILvarO, *P* > 0.999; Δ*epa_var* versus pILvarO, *P* < 0.0001. In both (**c**) and (**d**), statistical comparisons were made by doing a Kruskal-Wallis test, followed by Dunn’s multiple comparisons test.

**Fig. S10.**
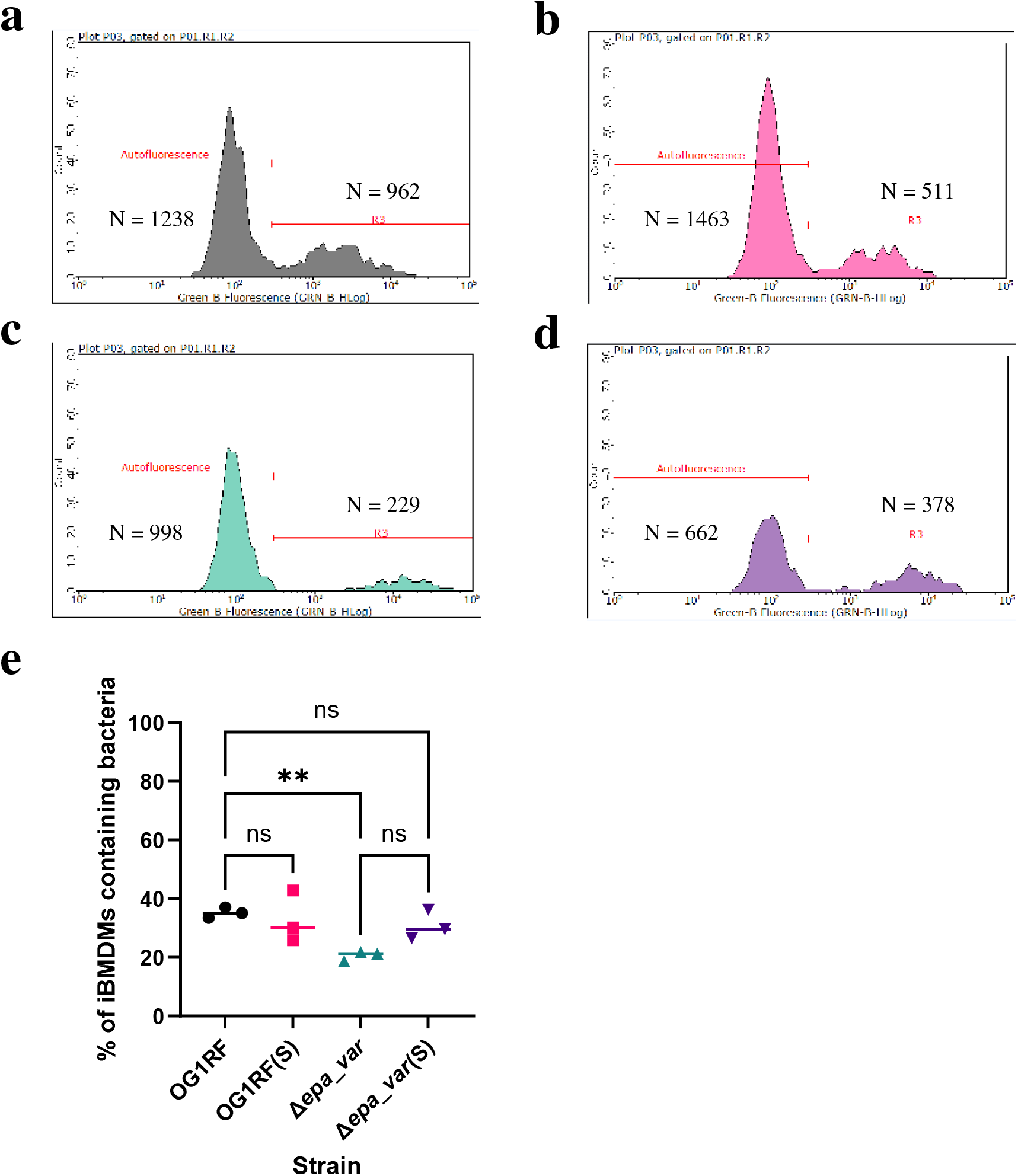
Phagocytosis of *E. faecalis* OG1RF or Δ*epa_var* with or without sonication beforehand. **(a-d)** Histograms plotting green fluorescence of iBMDMs following incubation with GFP-labelled OG1RF **(a)**, sonicated (S) OG1RF **(b)**, Δ*epa_var* **(c)**, or sonicated (S) Δ*epa_var* bacteria **(d)**. On each plot, the separation between bacteria-free (Autofluorescence) and bacteria-positive (gate R3) macrophages is indicated. Each plot represents one of three independent replicates performed for each treatment in this experiment. In this figure, N = number of iBMDMs within each gate. **(e)** Percentage of iBMDMs that did contain bacteria. Each value is the mean of three replicates per treatment. Statistical analysis was performed using a one-way ANOVA with Brown-Forsythe and Welch’s correction followed by Dunnett’s multiple comparisons test. *P*-values: OG1RF versus OG1RF(S), *P* = 0.995; OG1RF versus Δ*epa_var*, *P* = 0.0022; OG1RF versus Δ*epa_var*(S), *P* = 0.668; Δ*epa_var* versus Δ*epa_var*(S), *P* = 0.224. Key to *P*-values: ns, not significant; **, *P* < 0.01.

**Figure S11:**
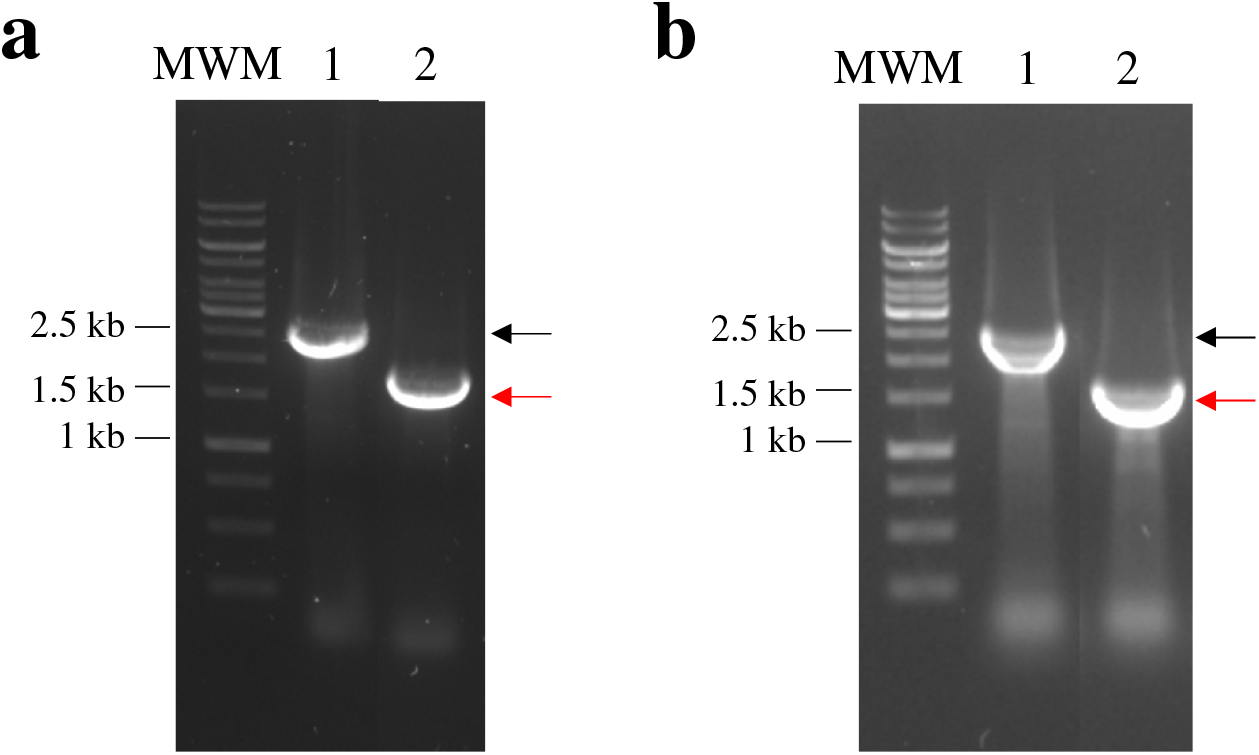
Gel electrophoresis of colony PCRs to characterize Δ*lgt* in-frame deletion mutants. Colony PCR using primers SM_0210 and SM_0211 was used to screen Δ*lgt* mutants in the OG1RF wild-type (**a**) and Δ*epa_var* backgrounds (**b**). The expected DNA band sizes corresponding to *lgt* (2,444 bp) and its deleted counterparts (Δ*lgt*, 1,643 bp) are indicated with black and red arrows, respectively. Lane 1, control colony PCR using OG1RF; lane 2, Δ*lgt* mutant.

**Figure S12:**
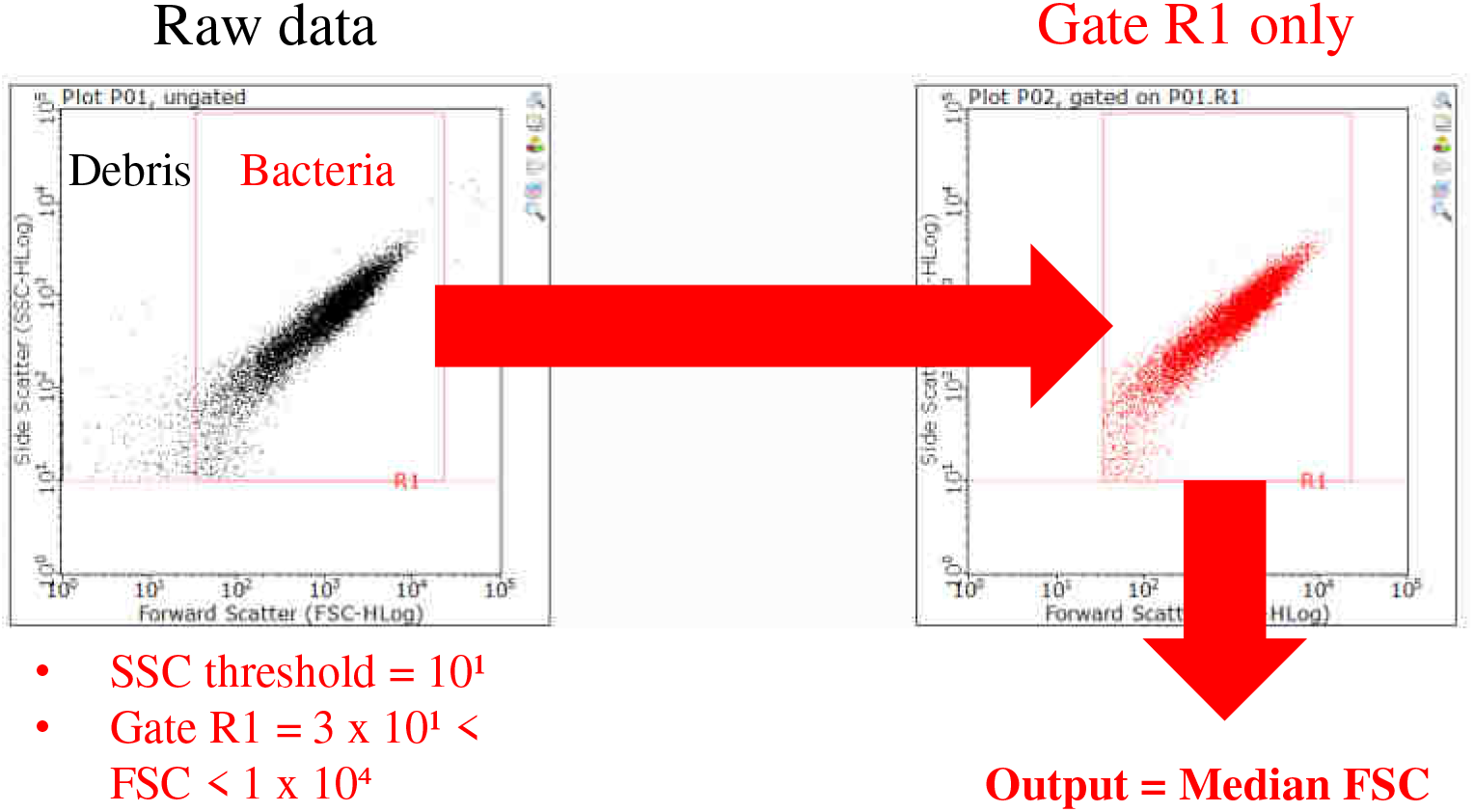
Flow cytometry gating strategy for *E. faecalis*. Data points were first plotted as a FSC log (x axis) versus SSC log scatter graph (left panel). Debris were excluded by (i) setting a threshold = 10^1^ for SSC values and (ii) drawing gate R1 spanning 3 x 10^1^ < FSC < 1 x 10^4^ (right panel). Median FSC was determined from all data points within gate R1. FSC, forward scatter; SSC, side scatter.

