## Supplementary tables for "Exploring the role of *E. faecalis* Enterococcal Polysaccharide Antigen (EPA) and lipoproteins in evasion of phagocytosis"

**Table S1: Bacterial strains, plasmids, and oligonucleotides.**

| Strains/plasmids/ oligonucleotides | Relevant properties/sequence [a] | Source |
|---|---|---|
| **Strains** | | |
| *Enterococcus faecalis* | | |
| OG1RF | Plasmid-free, virulent strain isolated from a human oral cavity | (1) |
| OG1RF (pGFP) | OG1RF derivative expressing GFP; Tet5 | (2) |
| OG1RF Δ*var* | OG1RF mutant with a complete deletion of the locus encoding EPA decorations including *OG1RF_11720* to *OG1RF_11706* | P. Serror |
| OG1RF Δ*var* (pGFP) | OG1RF Δ*var* derivative expressing GFP; Tet5 | This work |
| OG1RF Δ*var* (pIL252) | OG1RF Δ*var* derivative harbouring pIL252; Erm3 | P. Serror |
| OG1RF Δ*var* (pIL252, pGFP) | OG1RF Δ*var* (pIL252) derivative expressing GFP; Erm3, Tet5 | This work |
| OG1RF Δ*var* (pIL-O) | OG1RF Δ*var* derivative transformed with pIL-O encoding the EPA decoration locus from *E. faecalis* OG1RF | P. Serror |
| OG1RF Δ*var* (pIL-O, pGFP) | OG1RF Δ*var* (pIL-O) derivative expressing GFP; Erm3, Tet5 | This work |
| OG1RF Δ*var* (pIL-V) | OG1RF Δ*var* derivative complemented with pIL-V encoding the EPA decoration locus from *E. faecalis* V583 | P. Serror |
| OG1RF Δ*var* (pIL-V, pGFP) | OG1RF Δ*var* (pIL-V) derivative expressing GFP; Erm3, Tet5 | This work |
| OG1RF Δ*lgt* | OG1RF mutant with an in-frame deletion in *lgt* | This work |
| OG1RF Δ*lgt* (pGFP) | OG1RF Δ*lgt* derivative expressing GFP; Tet5 | This work |
| OG1RF Δ*lgt* (plgt) | OG1RF Δ*lgt* mutant complemented with a plasmid-encoded inducible *lgt* gene | This work |
| OG1RF Δ*lgt* (plgt, pGFP) | OG1RF Δ*lgt* (pTet_lgt) derivative expressing GFP; Erm30, Tet5 | This work |
| OG1RF Δ*var*Δ*lgt* | OG1RF mutant with an in-frame deletion in *lg* | This work |
| OG1RF Δ*var*Δ*lgt* (pGFP) | OG1RF Δ*var* derivative with an in-frame deletion in *lgt* expressing GFP; Tet5 | This work |
| OG1RF Δ*var*Δ*lgt* (plgt) | OG1RF Δ*var*Δ*lgt* mutant complemented with a plasmid-encoded inducible *lgt* gene; Erm30 | This work |
| OG1RF Δ*var*Δ*lgt* (plgt, pGFP) | OG1RF Δ*var*Δ*lgt* (plgt) derivative expressing GFP; Erm30, Tet5 | This work |
| V583* (VE18379) | V583 plasmidless derivative | (3) |
| V583* (pGFP) | V583* derivative expressing GFP; Tet5 | This work |
| V583* Δ*var* (VE18395) | V583* mutant with a complete deletion of the locus encoding EPA decorations including *EF2176* to *EF2164* | (3) |
| V583* Δ*var* (pIL252) (VE18927) | V583* Δ*var* derivative harbouring pIL252; Erm3 | (3) |
| V583* Δ*var* (pIL252, pGFP) | V583* Δ*var* (pIL252) derivative expressing GFP; Erm3, Tet5 | This work |
| V583* Δ*var* (pIL-V) (VE18930) | V583* Δ*var* derivative complemented with pILvarV encoding the EPA decoration locus from *E. faecalis* V583; Erm3 | (3) |
| V583* Δ*var* (pIL-V, pGFP) | V583* Δ*var* (pILvarV) derivative expressing GFP; Erm3, Tet5 | This work |
| V583* Δ*var* (pIL-O) | V583* Δ*var* derivative complemented with pILvarO encoding the EPA decoration locus from *E. faecalis* OG1RF; Erm3 | P. Serror |
| V583* Δ*var* (pIL-O pGFP) | V583* Δ*var* (pILvarO) derivative expressing GFP; Erm3, Tet5 | This work |
| *Escherichia coli* | | |
| TG1 (*repA*$^+$) | TG1 derivative encoding RepA for pGhost9 propagation at 37°C | (4) |
| NEB5α | Cloning strain | NEB |
| **Plasmids** | | |
| pGhost9 | Temperature-sensitive plasmid for gene replacement; ErmR | (5) |
| pG_lgt | pGhost9 derivative used to create an in-frame *lgt* deletion; ErmR | This work |
| pTetH | pAT18 derivative encoding TetR for tetracycline-inducible expression in *E. faecalis* | (6) |

| | | |
|---|---|---|
| pTetH_lgt | pTetH derivative used to complement the *lgt* mutation | This work |
| pMV158_GFP | pMV158 derivative consitutively expressing the *gfp* | (7) |
| pIL252 | Low-copy number plasmid used for epa complementation experiements; ErmR | (8) |
| pILvarO | pIL252 derivative encoding the *epa variable* locus from *E. faecalis* OG1RF (from the 8 last codons of *epaR* to the first 6 codons of *glpQ*) | P. Serror |
| pILvarV (pVE14388) | pIL252 derivative encoding the complete *epa variable* locus from *E. faecalis* V583 | (3) |
| **Oligonucleotides** | | |
| SM_0100 (pTetH_F) | GCTTGATCGTAGCGTTAACAGATCTACTC | |
| SM_0101 (pTetH_R) | CAAATTGTGGATGTGACCATGCGG | |
| SM_0171 (pGhost_F) | GTCACGACGTTGTAAAACGACGG | |
| SM_0172 (pGhost_R) | CTAGCGGACTCTAGAGGATCCCA | |
| SM_0194 (pG_lgt_H11) | TATAGGGCGAATTGGGTACCGGGCCCCCCCTCGAGTTTATTAGTGATGCGCCGTTTATGTGC | |
| SM_0195 (pG_lgt_H12) | TACTTGAGCTAACATCATCACATCTTCTCC | |
| SM_0196 (pG_lgt_H21) | GATGTGATGATGTTAGCTCAAGTAAAAATAACAACTTCTTAATCAACTTGATCAAAGAATATGG | |
| SM_0197 (pG_lgt_H22) | CGGACTCTAGAGGATCCCACCGCGGTGGCGGCCGCGCTAAACCACGAGTCATAATTGCCG | |
| SM_0210 (pG_lgt_H110) | TGACCGTTTACAATTATTTGGGAGC | |
| SM_0211 (pG_lgt_H220) | TTAAATCCCCAACACCACTTAAACC | |
| SM_0401 (pTetH_lgt_F) | TCATTGATAGAGTGAGCTCAAGGAGGAGACTGACCATGGAATTAGCTCAAGTAAATTCAATTGC | |
| SM_0402 (pTetH_lgt_R) | CTTTAGTGATGATGGTGATGGTGATGGTGGATCCAGAAGTTGTTATTTTTTTCTTTTGTTTCTCCCG | |

[a] Erm3, Erythromycin 3µgml$^{-1}$; Erm30, Erythromycin 30µg ml$^{-1}$; Tet5, Tetracycline 5µg ml$^{-1}$

**Table S2: Antibiotics used in this study.**

| Antibiotic | Stock concentration (mg/ml) | Working concentration (µg/ml) | |
|---|---|---|---|
| | | *E. faecalis* | *E. coli* |
| Ampicillin (Amp) | 100 | N/A | 100 |
| Erythromycin (Erm) | 30 | 30 or 3[a] | 200 |
| Gentamicin (Gen) | 50 | 250 | N/A |
| Tetracycline (Tet) | 10 | 5 | N/A |
| Vancomycin (Van) | 50 | 20 | **N/A** |

[a] Erm concentration is 30µg $ml^{-1}$ for pGhost and 3µg $ml^{-1}$ for pIL252; N/A, not applicable

**Table S3: Wavelengths and filters used for microscopy experiments.**

| Fluorophore | Excitation/emission wavelengths (nm) | Filter |
|---|---|---|
| AlexaFluor$^{TM}$ 555 NHS [a] ester | 555/572 | Texas Red |
| GFP [b] | 488/510 | FITC [c] |
| HADA [d] | 395/450 | DAPI [e] |
| Propidium iodide | 555/617 | Texas Red |

[a] NHS, *N*-hydroxy succinimide
[b] GFP, green fluorescent protein
[c] FITC, Fluorescein isothiocyanate
[d] HADA, hydroxycoumarin-carbonyl-amino-D-alanine
[e] DAPI, 4′,6-diamidino-2-phenylindole
